# A critical role for Dna2 at unwound telomeres

**DOI:** 10.1101/263228

**Authors:** Marta Markiewicz-Potoczny, Michael Lisby, David Lydall

## Abstract

Dna2 is a nuclease and helicase that functions redundantly with other proteins in Okazaki fragment processing, double strand break (DSB) resection and checkpoint kinase activation. Dna2 is an essential enzyme, required for yeast and mammalian cell viability. Here we report that numerous mutations affecting the DNA damage checkpoint suppress *dna2*Δ lethality in *Saccharomyces cerevisiae*. *dna2*Δ cells are also suppressed by deletion of helicases, *PIF1* and *MPH1*, and by deletion of *POL32*, a subunit of DNA polymerase δ. All *dna2*Δ cells are temperature sensitive, have telomere length defects, and low levels of telomeric 3’ single stranded DNA (ssDNA). Interestingly, Rfa1, a subunit of the major ssDNA binding protein RPA, and the telomere specific ssDNA binding protein Cdc13, often co-localize in *dna2*Δ cells. This suggests that telomeric defects often occur in *dna2*Δ cells. There are several plausible explanations for why the most critical function of Dna2 is at telomeres. Telomeres modulate the DNA damage response (DDR) at chromosome ends, inhibiting resection, ligation and cell cycle arrest. We suggest that Dna2 nuclease activity contributes to modulating the DNA damage response at telomeres by removing telomeric C-rich ssDNA and thus preventing checkpoint activation.

## Introduction

Dna2 is a conserved nuclease/helicase affecting 5’ processing of Okazaki fragments during lagging strand replication (Budd and Campbell, 1997), resection of DSBs/uncapped telomeres (Ngo et al., 2014), activation of DNA damage checkpoint pathways (Kumar and Burgers, 2013), resolution of G quadruplexes (Lin et al., 2013) and mitochondrial function (Budd et al., 2006, Duxin et al., 2009). Increased expression of *DNA2* is found in a broad spectrum of cancers, including leukemia, melanoma, breast, ovarian, prostate, pancreatic and colon cancers (Peng et al., 2012, Dominguez-Valentin et al., 2013, Strauss et al., 2014, Kumar et al., 2017, Jia et al., 2017, COSMIC). Dna2 is an important enzyme since its loss is lethal in human cell lines, mice, *C. elegans*, budding yeast and fission yeast (Budd et al., 1995, Kang et al., 2000, Lin et al., 2013). The amount of Dna2 in cells also seems to be important since *dna2*Δ/*DNA2* heterozygous mice show increased levels of aneuploidy-associated cancers and cells from these mice contain high numbers of anaphase bridges and dysfunctional telomeres (Lin et al., 2013).

In budding yeast Dna2 functions redundantly with other proteins in its various roles, and intriguingly, unlike Dna2, most of these proteins are not essential. For example, Rad27, Rnh201, Exo1 are all non-essential and are also involved in processing of 5’ ends of Okazaki fragments (Bae et al., 2001, Kao and Bambara, 2003). Exo1, Sgs1, Sae2, Mre11, Rad50, Xrs2, all non-essential, are involved in DSB resection (Mimitou and Symington, 2008, Zhu et al., 2008, Shim et al., 2010). Ddc1 (non-essential) and Dpb11 (essential) are involved in Mec1 (essential) checkpoint kinase activation (Navadgi-Patil and Burgers, 2009b, Puddu et al., 2008, Navadgi-Patil and Burgers, 2009a, Kumar and Burgers, 2013). Given that Dna2 often functions redundantly with non-essential proteins it is unclear what specific function or functions of Dna2 is/are so critical for cell viability.

A number of genetic and biochemical experiments had suggested that the most critical function of Dna2 is in processing long flaps at a small subset of 5’ ends of Okazaki fragments (Budd et al., 2011, Balakrishnan and Bambara, 2013). Dna2, is unique in that unlike the other 5’ nucleases (Rad27, Exo1, Rnh201), it can cleave RPA-coated single stranded DNA (ssDNA) (Stewart et al., 2008, Cejka et al., 2010, Levikova et al., 2013, Levikova and Cejka, 2015, Myler et al., 2016). RPA, the major eukaryotic ssDNA binding protein, binds ssDNA of 20 bases or more (Sugiyama et al., 1997, Rossi and Bambara, 2006, Balakrishnan and Bambara, 2013). Furthermore, RPA-coated ssDNA is potentially lethal because it stimulates DNA damage checkpoint responses (Lee et al., 1998, Zou and Elledge, 2003).

The two reported null suppressors of *dna2*Δ lethality, *rad9*Δ and *pif1*Δ, delete proteins that interact with RPA-coated ssDNA (Budd et al., 2006, Budd et al., 2011). Rad9 is important for the checkpoint pathway stimulated by RPA-coated ssDNA (Lydall, Weinert 1995). Pif1, a 5’ to 3’ helicase, increases the length of 5’ ssDNA flaps on Okazaki fragments, creating substrates for RPA binding, therefore checkpoint activation, and Dna2 cleavage (Pike et al., 2009, Levikova and Cejka, 2015). These genetic and biochemical data supported a model in which Dna2 is critical for cleaving RPA-coated long flaps from a subset of Okazaki fragments (Budd et al., 2011). However, more recently it was reported that other checkpoint mutations (*ddc1*Δ or *mec1*Δ), also affecting the response to RPA-coated ssDNA, did not suppress *dna2*Δ. It was suggested that specific interactions between Rad9 and Dna2 were important for the viability of *dna2*Δ *rad9*Δ cells, rather than the response to RPA-coated ssDNA per se (Kumar and Burgers, 2013).

In budding yeast, checkpoint mutations, such as *rad9*Δ and *ddc1*Δ, exacerbate fitness defects caused by general DNA replication defects (e.g. defects in DNA ligase, Pol α, Pol δ or Pol ɛ) (Weinert et al., 1994, Dubarry et al., 2015), but suppress defects caused by mutations affecting telomere function (e.g. defects in Cdc13, Stn1, Yku70) (Addinall et al., 2008, Holstein et al., 2017). The opposing effects of checkpoint mutations in general DNA replication or telomere-defective contexts is most likely explained by damage to non-coding telomeric DNA being comparatively benign in comparison to damage to coding DNA in the middle of chromosomes. By this logic the suppression of *dna2*Δ by *rad9*Δ implies that *dna2*Δ might cause telomere-specific rather than general chromosome replication defects. Furthermore, Dna2 localises to human and yeast telomeres (Choe et al., 2002, Chai et al., 2013, Lin et al., 2013), and *pif1*Δ, which suppresses *dna2*Δ, affects a helicase that is active at telomeres and affects telomere length (Dewar and Lydall, 2010, Budd and Campbell, 2013, Lin et al., 2013, Phillips et al., 2015). Thus, several lines of evidence suggest that Dna2 might play critical function(s) at telomeres.

To further explore whether Dna2 is important at telomeres we set out to clarify the effects of checkpoint pathways on fitness of *dna2*Δ mutants. We find that deletion of numerous DNA damage checkpoint mutations, all affecting responses to RPA-coated ssDNA, as well as deletions of Pif1 and Mph1 helicases, and Pol32, a subunit of Pol δ, suppress *dna2*Δ to a similar extent. These findings, along with a number of other telomere phenotypes lead us to suggest that the most critical function of Dna2 for cell viability is at telomeres. There are three possible substrates for Dna2 activity at telomeres: unwound telomeres, long flaps on terminal telomeric Okazaki fragments, and G4 quadruplexes formed on the G-rich ssDNA. We propose that Dna2 has its critical function in removing of RPA-coated, 5’ C-rich, ssDNA at telomeres.

## Results

### *dna2*Δ lethality is suppressed by checkpoint inactivation

To clarify the effect of DNA damage checkpoint gene deletions in *dna2*Δ cells, heterozygous *dna2*Δ *checkpoint*Δ diploid strains were sporulated, tetrads dissected and viable genotypes determined. We examined the effects of *RAD9*, *DDC1* and *MEC1*, affecting a checkpoint mediator protein, a component of the 9-1-1 checkpoint sliding clamp, and the central checkpoint kinase (homologue of human ATR), respectively, and all previously studied in the context of *dna2*Δ (Budd et al., 2011, Kumar and Burgers, 2013). We also examined *RAD17*, encoding a partner of Ddc1 in the checkpoint sliding clamp, *CHK1*, encoding a downstream checkpoint kinase, *RAD53*, a parallel downstream kinase, and *TEL1*, encoding the homologue of human ATM. As a positive control for suppression we also examined the effects of *PIF1*, encoding a 5’ to 3’ helicase, since *pif1*Δ, like *rad9*Δ, suppresses *dna2*Δ (Budd et al., 2006).

*dna2*Δ *rad9*Δ and *dna2*Δ *pif1*Δ strains are temperature sensitive (Budd et al., 2006, Budd et al., 2011) and therefore spores were germinated at 20°C, 23°C and 30°C to allow comparison of *dna2*Δ suppression frequencies at different temperatures. Interestingly, the effects of *rad9*Δ, *ddc1*Δ, *rad17*Δ, *chk1*Δ, and *mec1*Δ were very similar, they each permitted *dna2*Δ strains to form colonies at 20°C and 23°C but not at 30°C (Table 1, Figure 1, Figure S1a). In comparison, *pif1*Δ suppressed *dna2*Δ with higher efficiency and at higher temperatures, and *pif1*Δ *dna2*Δ colonies on germination plates were larger than those permitted by checkpoint gene deletions (Figure 1, Figure S1a, Table 1). *tel1*Δ and *rad53*Δ did not suppress *dna2*Δ, presumably because they have different roles in the DNA damage response. We conclude that *rad9*Δ, *ddc1*Δ, *rad17*Δ, *chk1*Δ and *mec1*Δ, but not *rad53*Δ and *tel1*Δ checkpoint mutations, suppress inviability caused by *dna2*Δ. These data suggest that *dna2*Δ causes lethal Rad9, Rad17, Ddc1, Chk1 and Mec1 mediated cell cycle arrest. Given that checkpoint mutations suppress *dna2*Δ and telomere defects (*cdc13-1*, *yku70*Δ and *stn1-13*) (Addinall et al., 2008, Holstein et al., 2017) but enhance DNA replication defects (Weinert et al., 1994, Dubarry et al., 2015), the pattern of *dna2*Δ genetic interactions strongly suggests that *dna2*Δ cells contained telomere defects.

**Figure 1.**
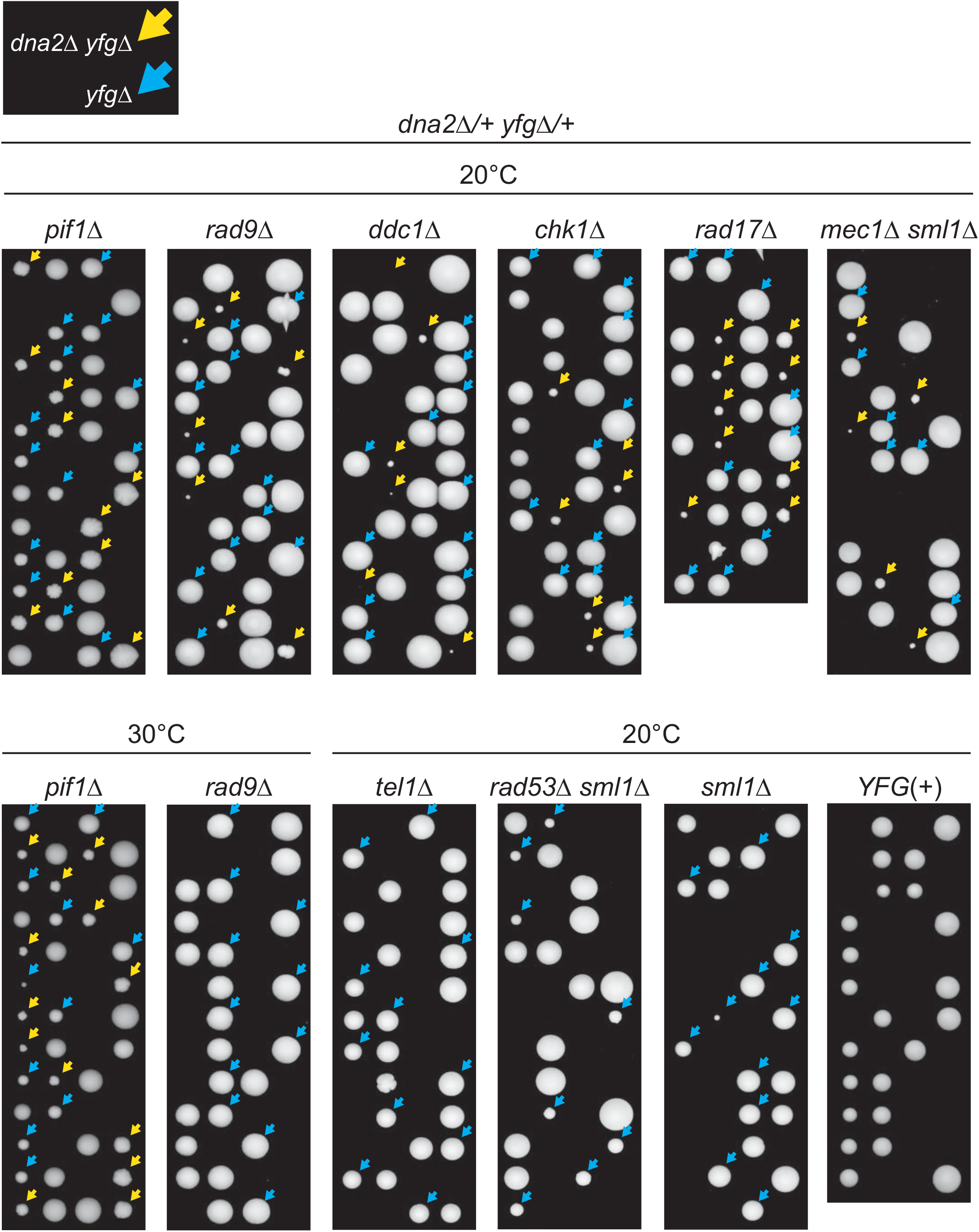
Checkpoint mutations permit growth of *dna2*Δ cells at 20°C. Diploids heterozygous for *dna2*Δ and *pif1*Δ, *rad9*Δ, *ddc1*Δ, *chk1*Δ, *rad17*Δ, *mec1*Δ *sml1*Δ, *tel1*Δ, *rad53*Δ *sml1*Δ or *sml1*Δ mutations were sporulated, tetrads dissected and spores germinated. Germination plates were incubated for 10-11 days at 20°C, or 3-4 days at 30°C. Strains of *dna2*Δ *yfg*Δ background are indicated by yellow arrows, and strains of *yfg*Δ background are indicated by blue arrows. Additional images of growth at 20°C, 23°C or 30°C are in Figure S1. Strains were: DDY1285, DDY874, DDY876, DDY878, DDY880, DDY958, DDY950, DDY947, DDY952, DDY1276, strain details are in Suppl. Table 1.

**Table 1.** *dna2*Δ suppression efficiency. 20°C, 23°C, 30°C – temp. at which spores were germinated. Left column is gene deleted in each *dna2*Δ/+ diploid. Viable *dna2*Δ *xyz*Δ-number of spores which germinated and formed visible colonies. Expected *dna2*Δ *xyz*Δ-expected number of viable *dna2*Δ *xyz*Δ strains if *xyz*Δ completely suppressed the *dna2*Δ inviable phenotype, based on the total number of tetrads dissected. E.g. 25% of *dna2*Δ/+ *rad9*Δ/+ spores should be *dna2*Δ *rad9*Δ, and 12.5% of *mec1*Δ/+ *sml1*Δ/+ *dna2*Δ/+ should be *mec1*Δ *sml1*Δ *dna2*Δ.

|  | 20°C |  | 23°C |  | 30°C |  |
| --- | --- | --- | --- | --- | --- | --- |
|  | Viable<br>*dna2*Δ<br>*xyz*Δ | Expected<br>*dna2*Δ<br>*xyz*Δ | Viable<br>*dna2*Δ<br>*xyz*Δ | Expected<br>*dna2*Δ<br>*xyz*Δ | Viable<br>*dna2*Δ<br>*xyz*Δ | Expected<br>*dna2*Δ<br>*xyz*Δ |
| XYZ | 0 | 12 |  |  | 0 | 12 |
| *rad9*Δ | 14 | 26 | 7 | 26 | 0 | 25 |
| *ddc1*Δ | 13 | 26 | 11 | 26 | 0 | 26 |
| *rad17*Δ | 20 | 23 | 12 | 26 | 0 | 25 |
| *chk1*Δ | 14 | 26 | 7 | 25 | 0 | 26 |
| *mec1*Δ *sml1*Δ | 16 | 49 |  |  | 0 | 12 |
| *pif1*Δ | 24 | 25 |  |  | 13 | 12 |
| *mph1*Δ | 10 | 26 |  |  | 0 | 13 |
| *pol32*Δ | 0 | 13 | 5 | 13 | 9 | 13 |
| *rad53*Δ *sml1*Δ | 0 | 19 |  |  | 0 | 25 |
| *tel1*Δ | 0 | 38 |  |  | 0 | 13 |
| *sml1*Δ | 0 | 13 |  |  |  |  |

### *DNA2* deletion causes temperature sensitivity

On germination plates *dna2*Δ *checkpoint*Δ colonies were often small and heterogeneous in size in comparison with *dna2*Δ *pif1*Δ colonies, implying that mutating checkpoint genes did not suppress the *dna2*Δ growth defects as efficiently as removing the Pif1 helicase (Figure 1). One explanation for this difference in colony size was that checkpoint mutations permitted only a limited number of cell divisions but that ultimately the *dna2*Δ *checkpoint*Δ double mutant clones would senesce and cease growth. To test this hypothesis, *dna2*Δ *checkpoint*Δ double mutants were passaged further. Interestingly, the opposite to senescence was observed, and *dna2*Δ *checkpoint*Δ mutants in fact became fitter and more homogeneous in colony size with passage and grew indefinitely (Figure 2a and Figure S2a). This suggests that *dna2*Δ *checkpoint*Δ double mutants originally grow quite poorly and that some type of adaptation to the absence of Dna2 occurs in *dna2*Δ *checkpoint*Δ mutants. We considered that additional suppressor mutations had arisen in *dna2*Δ *checkpoint*Δ mutants but backcross experiments did not support this hypothesis (Figure S1b). It was also clear that even different strains of the same genotype became similarly fit when passaged at 23°C, which is inconsistent with different suppressor mutations arising. However, all strains remained temperature sensitive for growth at higher temperatures, and growth at high temperature was more heterogeneous than growth at low temperature (Figure 2b, Figure S2b). Overall, passage of *dna2*Δ *checkpoint*Δ strains shows that they adapt to the absence of Dna2 but remain temperature sensitive for growth, presumably because ongoing cellular defects are more penetrant at higher temperature. Consistent with a previous study (Budd et al., 2006) *dna2*Δ *pif1*Δ strains, the least temperature sensitive genotype, formed smaller colonies at 36°C than at 30°C, showing that even these cells also have a temperature sensitive molecular defect (Figure 2b). We noted a similarity between *yku70*Δ and *dna2*Δ strains, since each genotype exhibits a temperature sensitive phenotype and is suppressed by checkpoint mutations (Maringele and Lydall, 2002). In the case of *yku70*Δ mutants high levels of 3’ ssDNA are generated at telomeres at high temperature (Maringele and Lydall, 2002).

**Figure 2.**
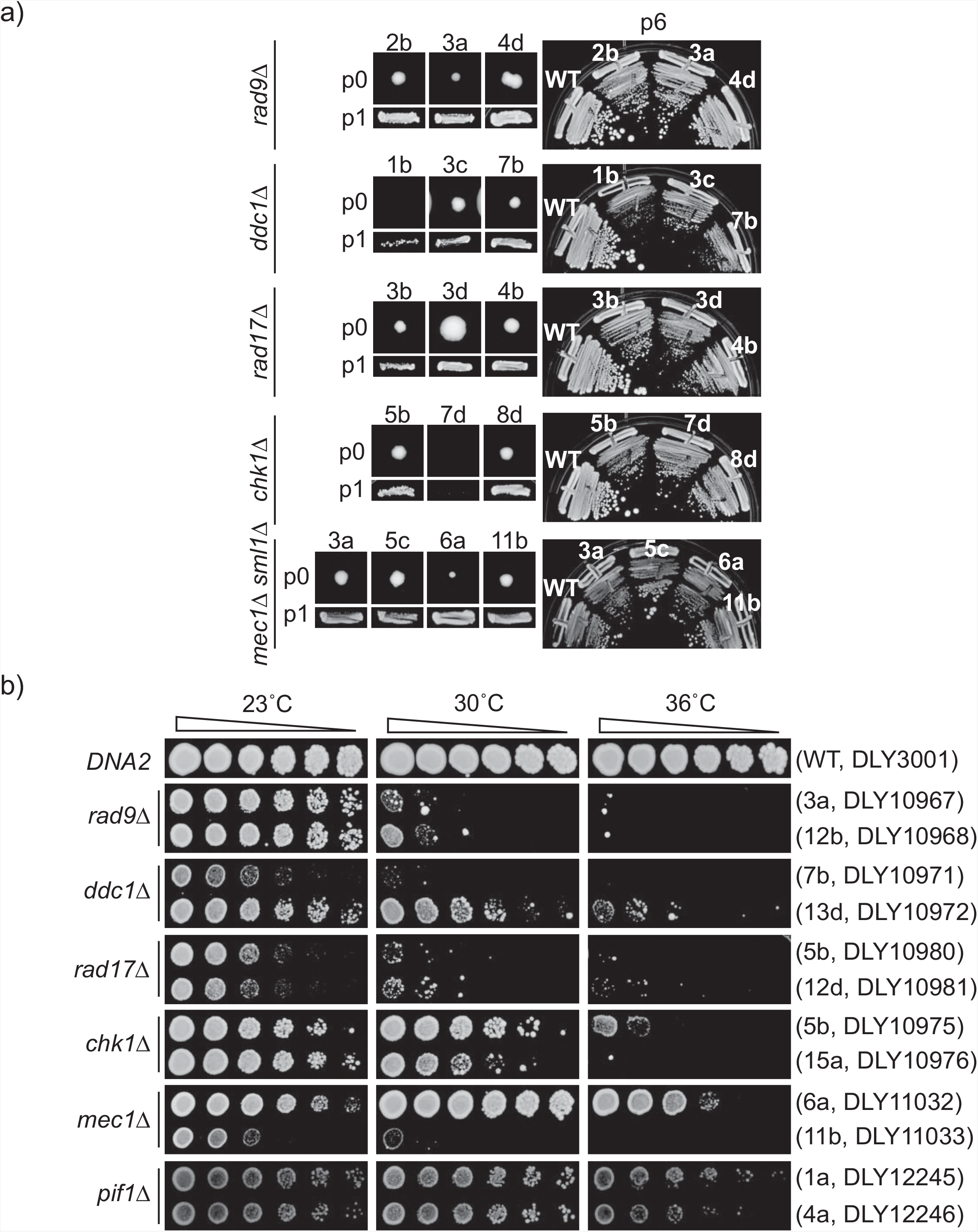
*dna2A* strains improve growth with passage, but remain temperature sensitive. a) Colonies of *dna2A yfgA* double mutants on germination plates (passage 0, p0) p1 (patched) and p6 (streaked) are shown.A single *DNA2* (WT) is used for comparison at p6. b) Spot test assays of strains at p6 (or p1 for *pif1A dna2A* strain). Strains of each genotype, at each temperature, were grown on single agar plates, but images have been cut and pasted to make comparisons easier. Original images are in Figure S2. Each colony position on germination plate from Figure 1 and strain numbers are indicated. Strain details are in Suppl. Table 1.

### *dna2*Δ cells have abnormal telomere length with limited ssDNA

We next tested whether Dna2 affects the structure of telomeric DNA. We first tested for increased levels of 3’ ssDNA at telomeres in *dna2*Δ cells, since this is seen in *yku70*Δ cells (Maringele and Lydall, 2002). Furthermore, in fission yeast Dna2 was shown to be involved in the generation of G-rich ssDNA at telomeres (Tomita et al., 2004). Importantly, it was reported that *dna2*Δ *rad9*Δ cells have abnormally low levels of telomeric 3’ G-rich ssDNA (Budd and Campbell, 2013). Consistent with what was reported for *rad9*Δ *dna2*Δ, *chk1*Δ *dna2*Δ, *mec1*Δ *dna2*Δ, *rad17*Δ *dna2*Δ, *ddc1*Δ *dna2*Δ and *pif1*Δ *dna2*Δ cells all showed low levels of 3’ G-rich ssDNA at telomeres in comparison with *DNA2* strains (Figure 3a-b, Figure S3, Figure S4). We conclude that all *dna2*Δ mutants have low levels of telomeric 3’ ssDNA. Interestingly, the *dna2*Δ ssDNA phenotype is opposite to that observed in other telomere defective strains (*cdc13-1* and *yku70*Δ mutants), which contain high levels of 3’ telomeric ssDNA (Maringele and Lydall, 2002). We also checked for 5’ C-rich ssDNA and saw no evidence for increased levels of telomeric C-rich ssDNA (Figure S5).

**Figure 3.**
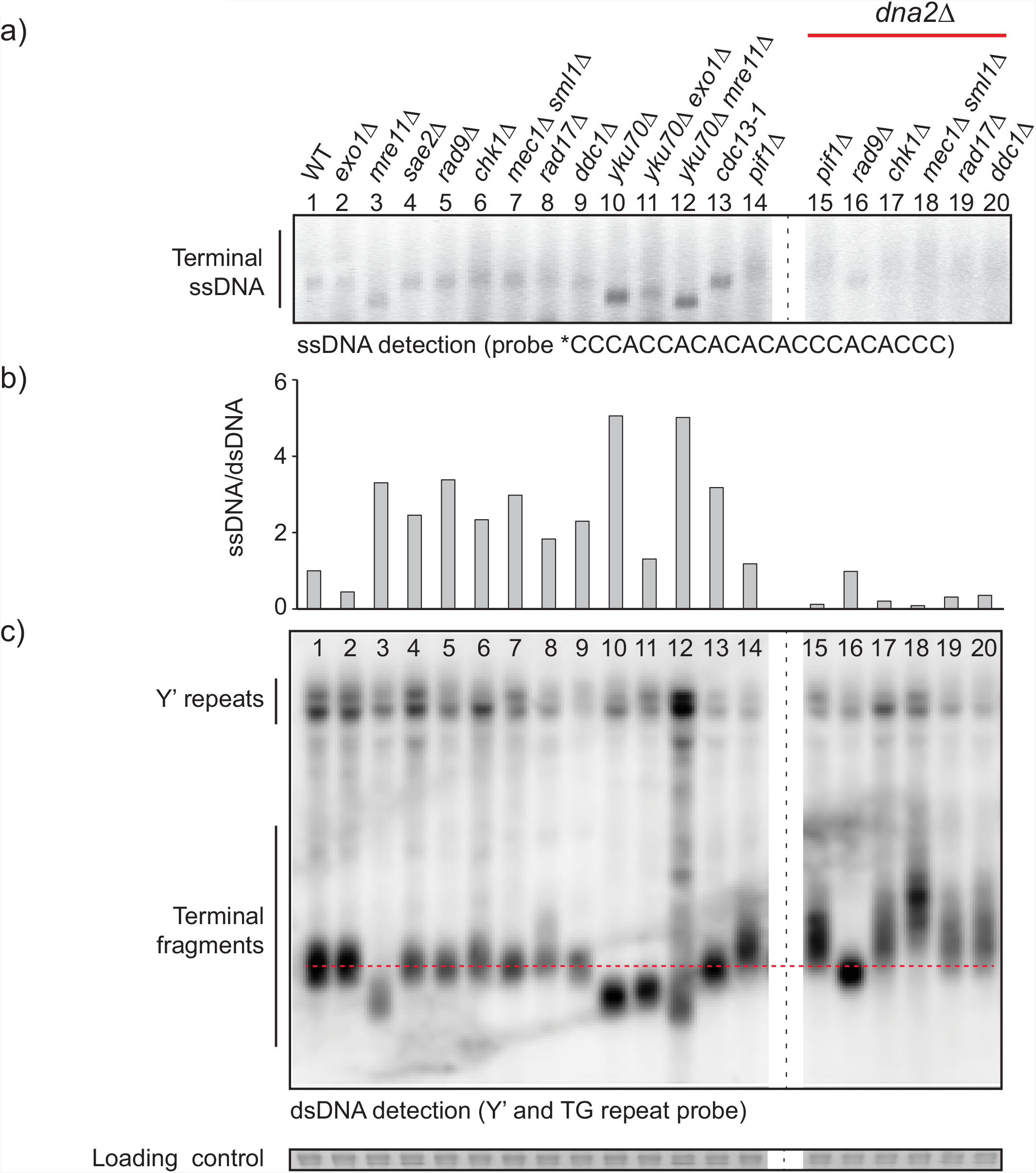
Telomeres of *dna2*Δ strains are abnormal and have low levels of ssDNA. a. An in-gel assay was performed to measure telomeric ssDNA. Saturated cultures were diluted 1:25 (*dna2*Δ strains) or 1:50 (other strains) and grown for 6 h until a concentration of approximately 107 cells/mL was attained. DNA was isolated from *dna2*Δ strains at passage 6, except for *dna2*Δ *pif1*Δ strain which is of unknown passage number. Strains were: WT (DLY3001), *exo1*Δ (DLY1272), *mre11*Δ (DLY4457), *sae2*Δ (DLY1577), *rad9*Δ (DLY9593), *chk1*Δ (DLY10537), *mec1*Δ *sml1*Δ (DLY1326), *rad17*Δ (DLY7177), *ddc1*Δ (DLY8530), *yku70*Δ (DLY6885), *yku70*Δ *exo1*Δ (DLY1408), *yku70*Δ *mre11*Δ (DLY1845), *cdc13-1* (DLY1108), *pif1*Δ (DLY4872), *pif1*Δ *dna2*Δ (DLY4690), *rad9*Δ *dna2*Δ (DLY10967), *chk1*Δ *dna2*Δ (DLY10975), *mec1*Δ *sml1*Δ *dna2*Δ (DLY11032), *rad17*Δ *dna2*Δ (DLY10981), *ddc1*Δ *dna2*Δ (DLY10973). Strain details are in Suppl. Table 1. * indicates a 5′ IRDye 800 label.
b. ssDNA and dsDNA were quantified using ImageJ analysis of the images shown in a) and c). The ratio of ssDNA/dsDNA is plotted and the wild type strain was given the value of “1”, all other ratios are expressed relative to the wild type. The telomeric regions quantified are indicated in Figure S3. Analysis of independent strains of the same genotypes is shown in Figure S4.
c. Southern blot was performed to measure telomeric dsDNA using a Y’-TG probe. SYBR Safe was used as a loading control, as previously described (Holstein et al., 2014).

**Figure 4.**
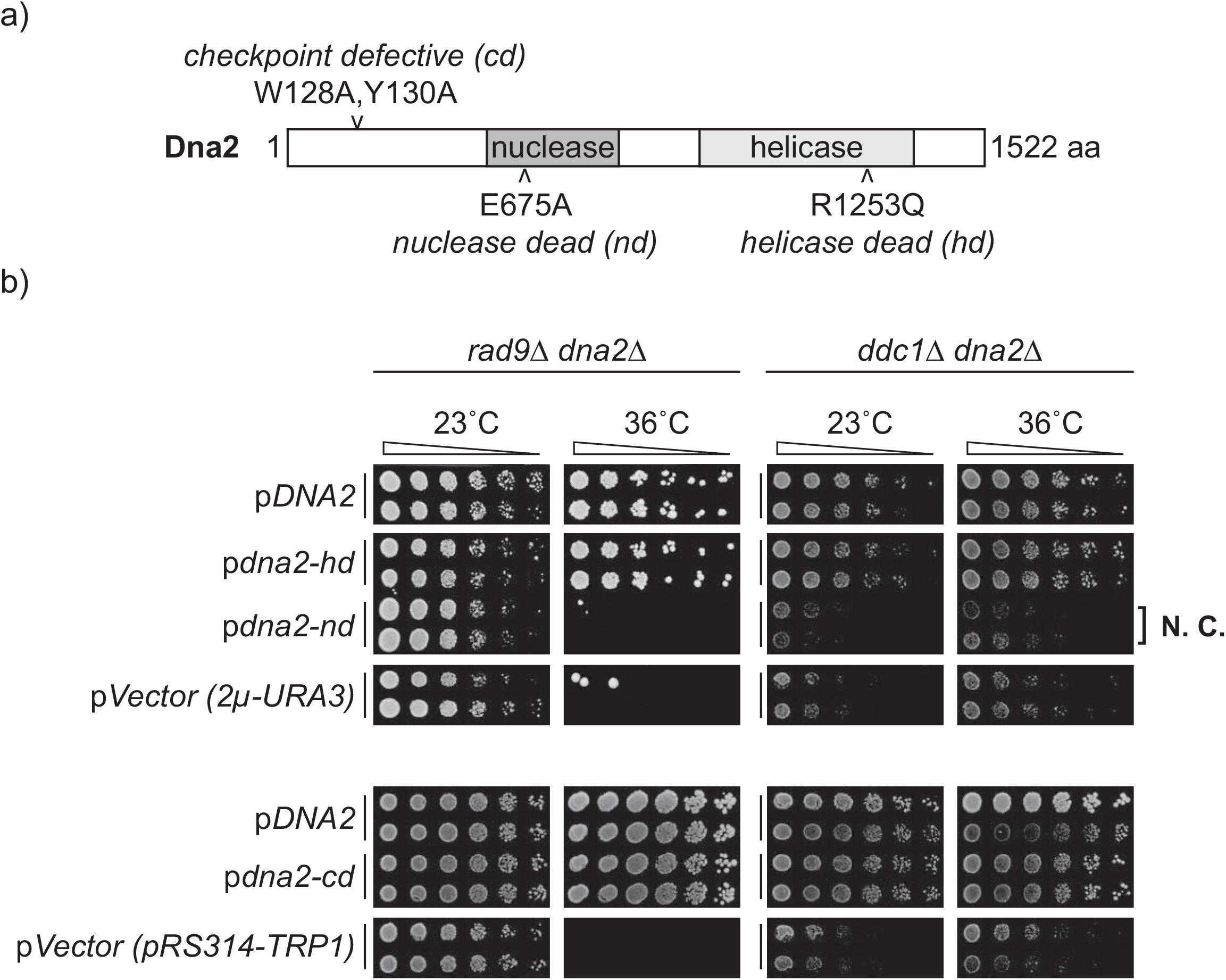
The nuclease domain of Dna2, but not helicase or checkpoint domains, confers viability of *dna2*Δ strains. a) Domain structure of yeast Dna2. Mutations affecting checkpoint, nuclease and helicase domains are indicated. b) Spot test assay performed as in Figure 2b. Strains from passage 6 of original colony 3a (*rad9*Δ *dna2*Δ, DLY10967), and 13d (*ddc1*Δ *dna2*Δ, DLY10973) were used for plasmid transformation. *rad9*Δ *dna2*Δ and *ddc1*Δ *dna2*Δ strains carrying DNA2, empty vector or helicase-dead, nuclease-dead or checkpoint-dead alleles of DNA2 were inoculated into 2 mL –URA or –TRP media for plasmid selection and cultured for 48 h, at 23°C. N.C. – no complementation. Original images are in Figure S8. Strain details are in Suppl. Table 1. Plasmid details are in Suppl. Table 2.

**Figure 5.**
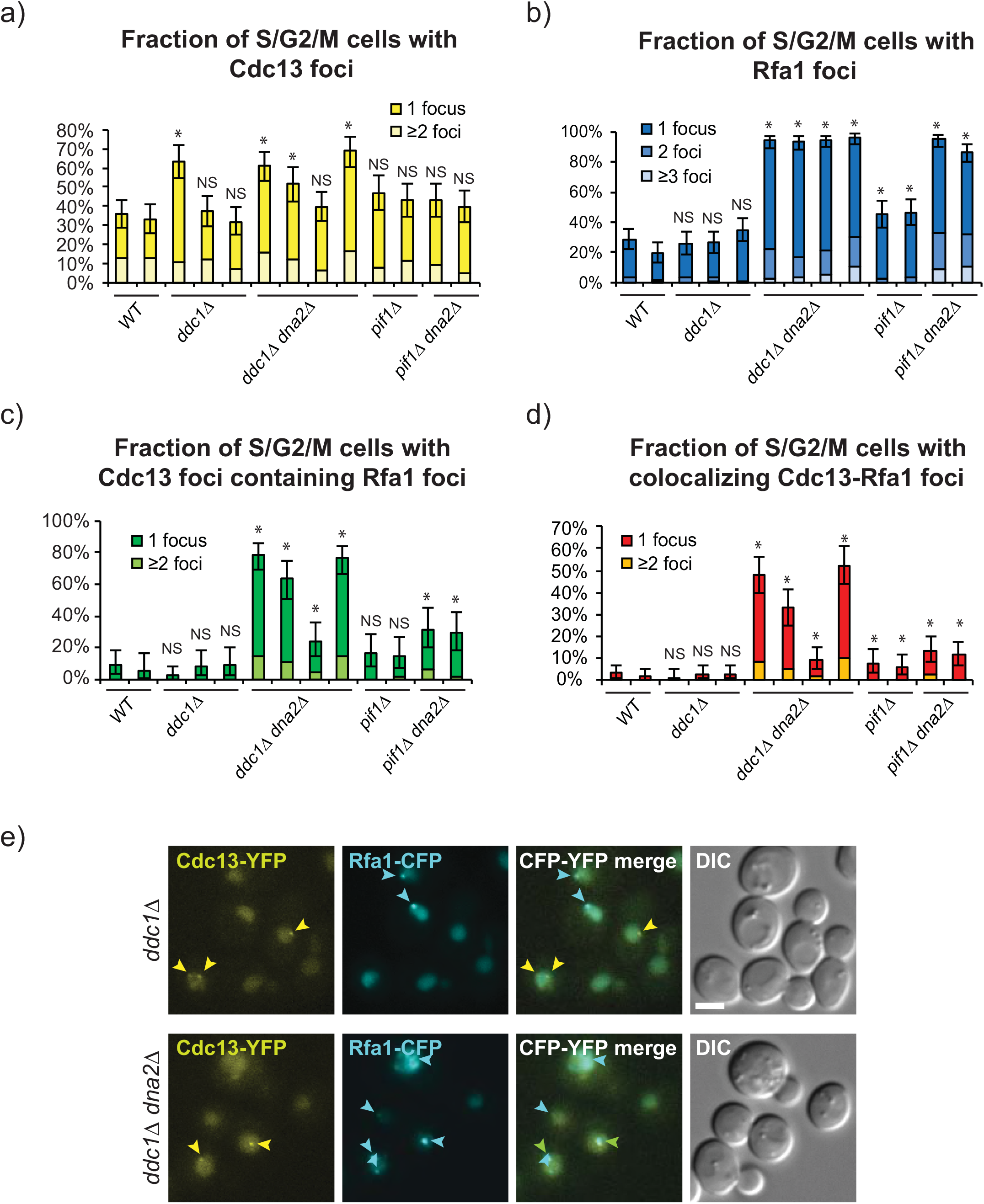
*dna2*Δ mutants accumulate CST and RPA, the ssDNA binding complexes. a-d) Percentages of Cdc13 foci, Rfa1 foci or colocalized Cdc13 –Rfa1 foci in *dna2*Δ and control strains are shown. a) Percentage of budded (S/G2/M) cells with either Cdc13 foci only or Cdc13-Rfa1 foci. b) Percentage of budded cells with either Rfa1 foci only or Cdc13-Rfa1 foci. c) Percentage of budded cells with Cdc13 foci that colocalize with Rfa1 foci. d) Percentage of budded cells with colocalizing Cdc13-Rfa1 foci. Error bars indicate 95 % confidence intervals (n=213-437, from two independent cultures of each strain). * - statistical significance (p<0.05) determined using Fisher’s exact test. NS - not significant. Strains are: WT (DLY12342, DLY12343), *ddc1*Δ (DLY12282, DLY12280, DLY12283), *ddc1*Δ *dna2*Δ (DLY12281, DLY12341, DLY12284, DLY12279), *pif1*Δ (DLY12346, DLY12347), *pif1*Δ *dna2*Δ (DLY12344, DLY12345). e) An example of live cell images is shown. Cdc13-Rfa1 co-localized foci are indicated by green arrow, Cdc13 foci by yellow arrows, and Rfa1 foci by blue arrows. Scale bar - 3 μm. Strain details are in Suppl. Table 1.

To search for other telomeric DNA phenotypes in *dna2*Δ strains we examined telomere length by Southern blot. Interestingly, the telomeres of *chk1*Δ *dna2*Δ, *mec1*Δ *dna2*Δ, *rad17*Δ *dna2*Δ, and *ddc1*Δ *dna2*Δ cells were long, and in fact longer and more diffuse, than *pif1*Δ strains, known to have very long telomeres (Schulz and Zakian, 1994) (Figure 3c, Figure S4, Figure S6). In contrast, and as reported before, *rad9*Δ *dna2*Δ telomeres were slightly shorter than the wild type length (Budd and Campbell, 2013). Rad9 is unique amongst checkpoint proteins because it binds chromatin and inhibits nuclease activity at telomeres and DSBs (Bonetti et al., 2015, Ngo and Lydall, 2015). Perhaps, therefore, the comparatively short telomere length in *rad9*Δ *dna2*Δ mutants reflects this chromatin binding function of Rad9 at telomeres. In summary, all *dna2*Δ mutants analysed have abnormal telomere lengths and low levels of 3’ G-rich ssDNA.

**Figure 6.**
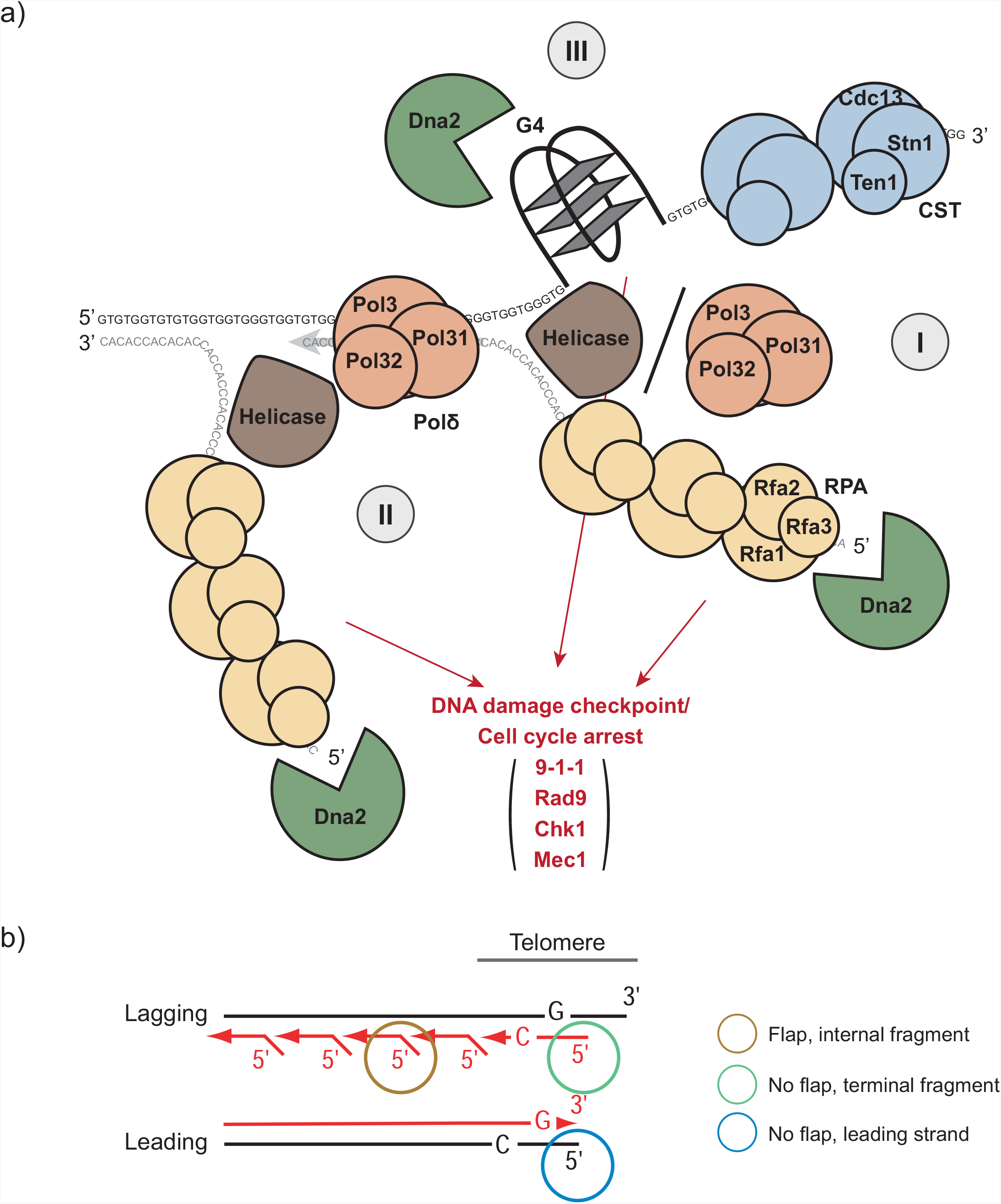
Three plausible roles for Dna2 in removing unwound RPA-coated ssDNA at telomeres. a) Three scenarios for Dna2 activity: I) 5‘ RPA-coated ssDNA cleavage at telomeric termini. Telomere ends are unwound by helicases, for example Pif1 or Mph1. The 3’ G-rich strand is bound by the CST, while the 5’ C-rich strand is bound by RPA, a substrate for Dna2 cleavage. II) Processing of long flaps on Okazaki fragments near telomeres. DNA polymerase δ displacement activity, stimulated by helicase(s), generates long flaps on an Okazaki fragment near telomere. Long C-rich flap, bound by RPA, are subjected to Dna2 cleavage. III) G-quadruplex unwinding and processing. G-quadruplexes formed on telomeric G-rich ssDNA are unwound or processed by Dna2. All proteins drawn to scale. b) Lagging and leading strand replication at telomeres. Short red arrows – Okazaki fragments on the lagging strand. Long red arrow – replicated leading strand. Brown circle – flap formed on internal Okazaki fragment. Green circle – no flap on the terminal telomeric Okazaki fragment. Blue circle – no flap on the leading strand template.

Long telomeres are present in telomerase deficient, recombination (*RAD52*) dependent survivors (Wellinger and Zakian, 2012). Recombination is also important to rescue stalled replication forks in telomeric sequences since the terminal location of telomeric DNA means that stalled forks cannot be rescued by forks arriving in the opposite direction, as elsewhere in the genome. Since the telomeres in *dna2*Δ strains were often long we wondered if recombination contributed to the viability of *dna2*Δ strains. Interestingly Rad52 did seem to contribute to the viability of *rad9*Δ *dna2*Δ and *ddc1*Δ *dna2*Δ strains (Figure S7). This strongly suggests that recombination dependent mechanisms help *dna2*Δ cells maintain viability.

### *Dna2* nuclease is critical in checkpoint-defective cells

Dna2 is a nuclease, a helicase and directly activates the central checkpoint kinase Mec1 (Kumar and Burgers, 2013). Any of these functions might be important at telomeres or elsewhere. To test which biochemical activity is most important to cell fitness we transformed nuclease, helicase, or checkpoint defective alleles of *DNA2* into *rad9*Δ *dna2*Δ or *ddc1*Δ *dna2*Δ cells, and measured growth at high temperature. It was clear that helicase dead and checkpoint defective alleles rescued the *dna2*Δ defect and permitted growth at high temperatures (Figure 4b, Figure S8). In contrast, the nuclease-defective allele of *DNA2* did not rescue the *dna2*Δ growth defect. We conclude that the most critical function of Dna2 in checkpoint defective yeast cells is its nuclease function.

### *dna2*Δ mutants contain RPA-bound telomeres

*dna2*Δ cells are temperature sensitive, have telomere length phenotypes and stimulate checkpoint pathways. However, paradoxically *dna2*Δ cells have reduced levels of telomeric ssDNA when measured by in-gel assay. We reasoned that one plausible function for Dna2 nuclease activity was removal of ssDNA present *in vivo* that was not detectable *in vitro*. That is, that unwound terminal telomeric DNA formed Y shaped structures *in vivo*, with splayed arms of G-rich and C-rich ssDNA. The 5’ C-rich and 3’ G-rich ssDNA should bind RPA and CST (Cdc13, Stn1 and Ten1) (Nugent et al., 1996), respectively, with the RPA-coated 5’ ssDNA stimulating DNA damage checkpoint pathways. The ssDNA present on the arms of Y shaped telomeres *in vivo* might not be detected by in-gel assays because complementary ssDNA strands would reanneal during DNA purification. Finally, telomere unwinding might be catalysed by helicases (for example, Pif1), and high temperature, explaining the effects of *pif1*Δ and temperature on fitness of *dna2*Δ cells.

Most eukaryotic cells contain 3’ ssDNA overhangs on the G-rich strand of telomeric DNA, and this ssDNA is bound by proteins such as Pot1 and CST. If unwound telomeres occur in *dna2*Δ cells, then CST should still bind the 3’ strand, but in addition RPA could bind the C-rich 5’ strand and stimulate the checkpoint. Presumably, in such case both RPA and CST complexes would co-localize at telomeres and presumably stop the stimulation of the checkpoint pathway. To explore RPA and CST localization the two largest subunits of each complex, Cdc13 and Rfa1, were tagged with YFP and CFP, respectively, and their localization in *dna2*Δ cells was examined by live cell microscopy.

We examined Cdc13 and Rfa1 foci in *ddc1*Δ *dna2*Δ, *pif1*Δ *dna2*Δ cells and WT, *ddc1*Δ, *pif1*Δ controls. Since some of these cells grew poorly, and may have altered cell cycle distributions, we counted foci in budded cells (S/G2/M) since this is when RPA foci are more likely to be present (Figure 5). We observed broadly similar fractions of cells with Cdc13 foci in all cultures at the level of 30-70%, but checkpoint defective strains, *ddc1*Δ and *ddc1*Δ *dna2*Δ, had somewhat higher levels (closer to 70%) (Figure 5a). In G1 cells the number of Cdc13 foci was smaller (less than 20%), but consistently *ddc1*Δ *dna2*Δ cells tended to have slightly higher levels (on average 15%) (Figure S9a). We conclude *DNA2* deletion has no strong effect on Cdc13 foci formation.

We also searched for Rfa1 foci and observed that on average 30% of budded and 10% of unbudded control cells contained Rfa1 foci (Figure 5b, Figure S9b). In contrast, *ddc1*Δ *dna2*Δ and *pif1*Δ *dna2*Δ cultures contained a much higher fraction of budded cells with Rfa1 foci. Generally, greater than 80% of *ddc1*Δ *dna2*Δ and *pif1*Δ *dna2*Δ cells, and approximately 40% of *pif1*Δ cells, contained at least one Rfa1 focus (Figure 5b), suggesting that high levels of DNA damage and ssDNA are present in these strains. In G1 cells the number of Rfa1 foci was smaller (up to 80%), and cells hardly ever contained more than one Rfa1 focus (Figure S9b).

If the Rfa1 foci observed in *dna2*Δ cells were primarily at telomeres, rather than at DSBs or long flap on Okazaki fragments elsewhere in the genome, then Rfa1 foci in *dna2*Δ cells should preferentially localize at telomeres. Assuming that Cdc13 foci are at telomeres (Khadaroo et al., 2009), then more than 60% of these telomeric loci in *ddc1*Δ *dna2*Δ budded cells co-localized with Rfa1 (Figure 5c). In contrast, less than 10% of Cdc13 foci contained Rfa1 in WT or *ddc1*Δ budded cells, suggesting little Rfa1 at telomeres in WT or *ddc1*Δ strains. This suggests that RPA bound ssDNA occurs at high frequency near telomeres in *ddc1*Δ *dna2*Δ cells. *pif1*Δ *dna2*Δ cells contained nearly as many Rfa1 foci and Cdc13 foci as *ddc1*Δ *dna2*Δ cells, but less Cdc13 foci contained Rfa1, approximately 30%. We conclude that *pif1*Δ *dna2*Δ cells have less RPA bound ssDNA at telomeres than *ddc1*Δ *dna2*Δ cells. Interestingly, *pif1*Δ single mutants also contained more Rfa1 foci than wild type cells, and more co-localization of Rfa1 and Cdc13, at approximately 5% (Figure 5b-d). This suggests that *pif1*Δ cells, which contain long telomeres, show comparatively high levels of RPA binding at telomeres, possibly due to the difficulty of replicating through long stretches of telomeric DNA.

Overall, of all the genotypes examined *ddc1*Δ *dna2*Δ mutants had the highest fraction of Cdc13 foci that contain Rfa1, Rfa1 foci that contain Cdc13, and Cdc13-Rfa1 foci (Figures 5c-d, Figure S9f). These data are consistent with a model in which both G-rich and C-rich ssDNA are found at high levels at telomeres in *ddc1*Δ *dna2*Δ cells. Interestingly, *pif1*Δ *dna2*Δ cells also contained increased levels of CST/RPA-bound ssDNA, suggesting that Pif-independent helicases may unwind telomeric C-rich and G-rich ssDNA in the absence of Pif1 to generate substrates for RPA binding.

### *dna2*Δ lethality is suppressed by *mph1*Δ and *pol32*Δ, but not *sgs1*Δ

To search for additional activities that might, like Pif1, unwind telomeric DNA we examined genes affecting likely candidates. Sgs1 was a candidate since it functions with Dna2 in resection of DSBs and uncapped telomeres (Cejka et al., 2010, Ngo et al., 2014), but its deletion did not suppress *dna2*Δ (Figure S10a), as has been reported by others (Hoopes et al., 2002, Weitao et al., 2003, Budd et al., 2005). On this basis Sgs1 does not seem to contribute to telomere unwinding, or if it does, it also has other functions which are essential in *dna2*Δ strains.

We examined Mph1, because like Pif1, Mph1 stimulates Dna2 activity on 5’ flaps *in vitro* (Kang et al., 2009). Interestingly, *mph1*Δ suppressed *dna2*Δ. The effect of *mph1*Δ was similar to checkpoint mutations, but not as strong as *pif1*Δ (Figure S10a-c). Therefore loss of Mph1, a 3’ to 5’ helicase, like loss Pif1, a 5’ to 3’ helicase, suppresses the inviability of *dna2*Δ cells. Given the polarity of the Mph1 helicase it would most likely engage with the 3’ G-rich overhanging strand to unwind telomeric DNA, and compete with CST for this substrate. To test this hypothesis, *mph1*Δ was combined with *cdc13-1* and the temperature-sensitive phenotype scored. Interestingly, *mph1*Δ mildly suppresses the temperature dependent growth defects of *cdc13-1* mutants (Figure S10d). This suggests that Mph1 and CST compete to bind the same G-rich strand at telomeres and is consistent with the idea that Mph1 engages with the 3’ telomeric overhang to unwind telomeric double stranded DNA (dsDNA).

Finally we tested Pol32, a DNA Pol δ sub-unit, which helps displace 5’ ends of Okazaki fragments. It had been reported that *pol32*Δ suppresses some alleles of *DNA2*, and to weakly suppress *dna2*Δ (Budd et al., 2006, Stith et al., 2008). Interestingly, we confirmed that *pol32*Δ suppressed *dna2*Δ. In contrast to checkpoint mutations *pol32*Δ suppressed *dna2*Δ at high temperature (30°C, and also 23°C) but not at 20°C (Figure S10a-c). This temperature dependent suppression may be explained by the fact that *pol32*Δ mutants are cold sensitive (Gerik et al., 1998).

## Discussion

We report that loss of proteins affecting numerous aspects of the DNA damage response permit budding yeast cells to divide indefinitely in the absence of the essential protein Dna2. Loss of DNA damage checkpoint proteins (Rad9, Ddc1, Rad17, Chk1 and Mec1), or Pif1, a 5’ to 3’ helicase, Mph1, a 3’ to 5’ helicase, or Pol32, a DNA polymerase δ subunit, suppress the inviability of *dna2*Δ cells. The suppression of *dna2*Δ by checkpoint mutations makes *dna2*Δ mutants more similar to telomere defective strains than general DNA replication defective strains (Dubarry et al., 2015). Consistent with this *dna2*Δ strains show telomere length phenotypes and a high-degree of co-localisation of Cdc13, a telomeric G-rich ssDNA binding protein, and Rfa1, a more general ssDNA binding protein *in vivo*. *dna2*Δ mutants are also temperature sensitive and have low levels of telomeric G-rich ssDNA. The nuclease function of Dna2, but not helicase and checkpoint functions, is critical to confer the viability of *dna2*Δ *checkpoint*Δ strains at high temperature.

The low levels of telomeric 3’ ssDNA that we detect at telomeres of *dna2*Δ mutants by *in vitro* in-gel assay, is the opposite phenotype to the high levels of 3’ ssDNA found at telomeres in other telomere defective strains suppressed by checkpoint gene mutations (for example, *cdc13-1* and *yku70*Δ mutants) (Maringele and Lydall, 2002, Ngo et al., 2014). Our explanation is that high levels of RPA-coated C-rich ssDNA and comparatively normal levels of CST-coated G-rich ssDNA are present at unwound telomeres of *dna2*Δ cells *in vivo*. This is detected as co-localization by live cell imaging but when DNA is extracted it renatures during purification and ssDNA is not detected.

There are at least three plausible scenarios for why Dna2 might have its most critical functions at, or near, telomeres (Figure 6a). One model, that best fits all our data, is that Dna2 nuclease activity removes potentially harmful, RPA-coated 5’ C-rich ssDNA, at the termini of telomeres (Figure 6a, scenario I). In this model helicases, like Pif1 or Mph1, unwind the telomeric termini. The G-rich strand is bound by the telomeric CST complex, and is presumably quite benign, but the C-rich strand is bound by RPA and potentially stimulates DNA damage checkpoint activity. Pol32, a subunit of DNA polymerase δ with strand displacement activity (Podust et al., 1995, Maga et al., 2001), might also generate ssDNA at the telomeric terminus, if CST recruits Pol α for lagging strand fill-in, which in turn recruits Pol δ (Waga and Stillman, 1998, Maga et al., 2000, Burgers, 2009).

Another potential role for Dna2 at telomeres is in removing long flaps of sub-telomeric Okazaki fragments (Figure 6a, scenario II). Finally, Dna2 nuclease activity may be needed at stalled replication forks in telomeric regions (Figure 6a, scenario III). For example, mammalian and yeast telomeres are G-rich, difficult to replicate and can form G quadruplexes that might be processed by Dna2 (Gilson and Geli, 2007, Masuda-Sasa et al., 2008, Lin et al., 2013, Maestroni et al., 2017). At other genomic locations other substrates for Dna2, e.g. DSBs, or stalled replication forks, can also occur (Ngo et al., 2014, Hu et al., 2012), but our evidence is that telomeres are particularly reliant on Dna2.

If Dna2 acts at the very termini of telomeres (Figure 6a, scenario I), either the lagging strand, the leading strand or both, might be targets for Dna2 (Figure 6b). It is well established that the leading and lagging strands of telomeres are processed by different mechanisms (Parenteau and Wellinger, 1999, Wu et al., 2012, Bonetti et al., 2013, Soudet et al., 2014). After lagging strand replication is complete the very terminus cannot be fully replicated because of the end replication problem. Irrespective of whether the most terminal Okazaki fragment is created by passage of the replication fork, or CST recruitment of Pol α, it is unusual, in that unlike more than 99% of the other Okazaki fragments, it will not contain a flap at its 5’ end (Figure 6b). Perhaps the absence of a flap, and/or a polymerase, facilitates helicase engagement. The leading strand telomere end, which is thought to be blunt after the replication fork has passed, may also be susceptible to helicase activities.

We and others (Budd and Campbell, 2013) have shown that *dna2*Δ *rad9*Δ cells have a short telomere phenotype. All other *dna2*Δ strains, including other checkpoint defective strains, have long telomeres. Hence it is not telomere length per se that determines the survival of *dna2*Δ cells. Rad9, like its human orthologue 53BP1, binds chromatin and inhibits resection at telomere defective *cdc13-1* cells and at DSBs (Iwabuchi et al., 2003, Lazzaro et al., 2008, Bunting et al., 2010, Ngo and Lydall, 2015). Perhaps Rad9 binding to chromatin also inhibits helicase activity, telomere unwinding and nuclease activity. Presumably unwound telomeres are also more susceptible to nucleases (other than Dna2). Consistent with this, the 9-1-1 complex recruits Dna2 and Exo1 nuclease to uncapped telomeres (Ngo and Lydall, 2015), and *ddc1*Δ *dna2*Δ and *rad17*Δ *dna2*Δ mutants, defective in 9-1-1, have long telomeres.

Telomeres in all organisms are difficult to replicate and need to be protected from the harmful aspects of the DNA damage response. Telomeric structures like t-loops, and proteins like CST, shelterin and the Ku heterodimer, may help protect telomeric DNA from being unwound by helicases. Our experiments in yeast suggest that Dna2 is critical for removing RPA-coated C-rich ssDNA at unwound telomeres. *DNA2* is an essential gene in budding and fission yeasts, *C. elegans*, mice and human cells. Interestingly, *C. elegans dna2*Δ mutants show temperature-dependent delayed lethality (Lee et al., 2003), suggesting that temperature-dependent telomere unwinding in *C. elegans* creates substrates for Dna2 nuclease activity at high temperatures.

Dna2 localizes at telomeres in yeast, humans and mice, and Dna2 affects telomere phenotypes in all these organisms (Choe et al., 2002, Lin et al., 2013). Dna2, checkpoint proteins, Pif1 and Mph1 helicases as well as Pol32 are all conserved between human and yeast cells, and affect telomere related human diseases, such as cancer, suggesting our observations may be relevant to human disease (Paeschke et al., 2013, Byrd and Raney, 2015, Ceccaldi et al., 2016). It will be interesting to see if telomere specific functions for Dna2 are conserved across eukaryotes.

## Materials and Methods

### Yeast culture and passage

All yeast strains were in W303 background and *RAD5*+ and *ade2-1*, except strains used for microscopy which were *ADE2* (Table S1). Media were prepared as described previously and standard genetic techniques were used to manipulate yeast strains (Sherman et al., 1986). YEPD (1L: 10 g yeast extract, 20 g bactopeptone, 50 mL 40 % dextrose, 15 mL 0.5 % adenine, 935 mL H20) medium was generally used. Dissected spores were germinated for 10-11 days at 20°C, 7 days at 23°C or 3-4 days at 30°C. Colonies from spores on germination plates were initially patched onto YEPD plates and grown for 3 days. Next these were streaked for single colonies and incubated for 3 days at 23°C. Thereafter, 5-10 colonies of each strain were pooled by toothpick and streaked for single colonies every three days.

### Yeast spot test assays

5-10 colonies were pooled, inoculated into 2 mL YEPD and grown to saturation on a wheel at 23°C. Saturated cultures were 5-fold serially diluted in sterile water (40 μL: 160 μL) in 96-well plates. Cultures were transferred onto rectangular YEPD agar plates with a rectangular pin tool, and incubated at the indicated temperatures for 3 days before photography, unless stated otherwise.

### In-gel assay/Southern blots

In-gel assays were performed as previously described (Dewar and Lydall, 2012) with minor modifications. Infrared 5’ IRDye 800 probes were used (AC probe: M3157, *CCCACCACACACACCCACACCC*; TG probe: M4462, *GGGTGTGGGTGTGTGTGGTGGG*, Integrated DNA Technologies). No RNAse was used during nucleic acid purification. Samples were run on a 1 % agarose gel in 0.5× TBE (50 V/ 3 h), and the probe was detected on a LI-COR (Odyssey) imaging system. ssDNA was quantified using ImageJ. The gel was then placed back in an electrophoresis tank, run for 2 more hours, and processed for Southern blot. Then gel was stained using SYBR Safe, and DNA was detected using a Syngene’s G:BOX imaging system. DNA was then transferred to a positively charged nylon membrane. The membrane was hybridized with a 1 kbp Y′ and TG probe as preivously described (Holstein et al., 2014). Loading controls were generated by foreshortening the full-sized SYBR Safe-stained gel images using Adobe Illustrator CS6.

### Yeast live cell imaging

Cells were grown shaking in liquid SC+Ade (synthetic complete medium supplemented with 100 μg/mL adenine) medium at 25°C to OD_600_ = 0.2−0.3 and processed for fluorescence microscopy as described previously (Silva et al., 2012). Rfa1 was tagged with cyan fluorescent protein (CFP, clone W7) (Heim and Tsien, 1996) and Cdc13 with yellow fluorescent protein (YFP, clone 10C) (Ormo et al., 1996, Khadaroo et al., 2009). Fluorophores were visualized with oil immersion on a widefield microscope (AxioImager Z1; Carl Zeiss) equipped with a 100× objective lens (Plan Apochromat, NA 1.4; Carl Zeiss), a cooled CCD camera (Orca-ER; Hamamatsu Photonics), differential interference contrast (DIC), and an illumination source (HXP120C; Carl Zeiss). Eleven optical sections with 0.4 μm spacing through the cell were imaged. Images were acquired and analysed using Volocity software (PerkinElmer). Images were pseudocoloured according to the approximate emission wavelength of the fluorophores.

## Acknowledgments

This work was funded by EU Marie Curie ITN network CodeAge (FP7-PEOPLE-2011-ITN) and the BBSRC (BB/M002314/1). The Danish Agency for Science, Technology and Innovation (DFF) and the Villum Foundation supported the work performed by ML. We are particularly grateful to Lata Balakrishnan, Peter Burgers, Judy Campbell, Laura Maringele, and Duncan Smith for advice and input.

---

**Supplementary Table 1.**
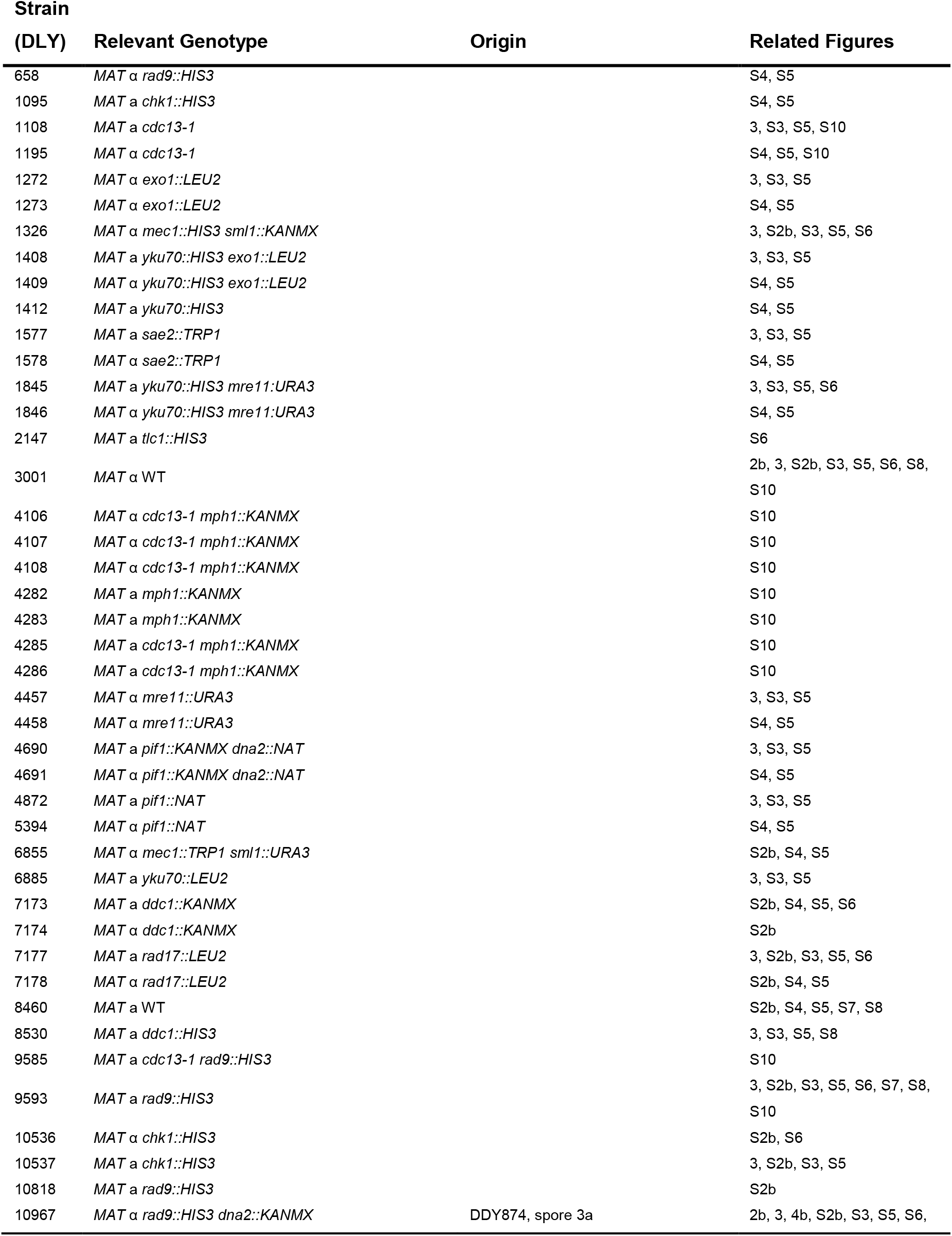
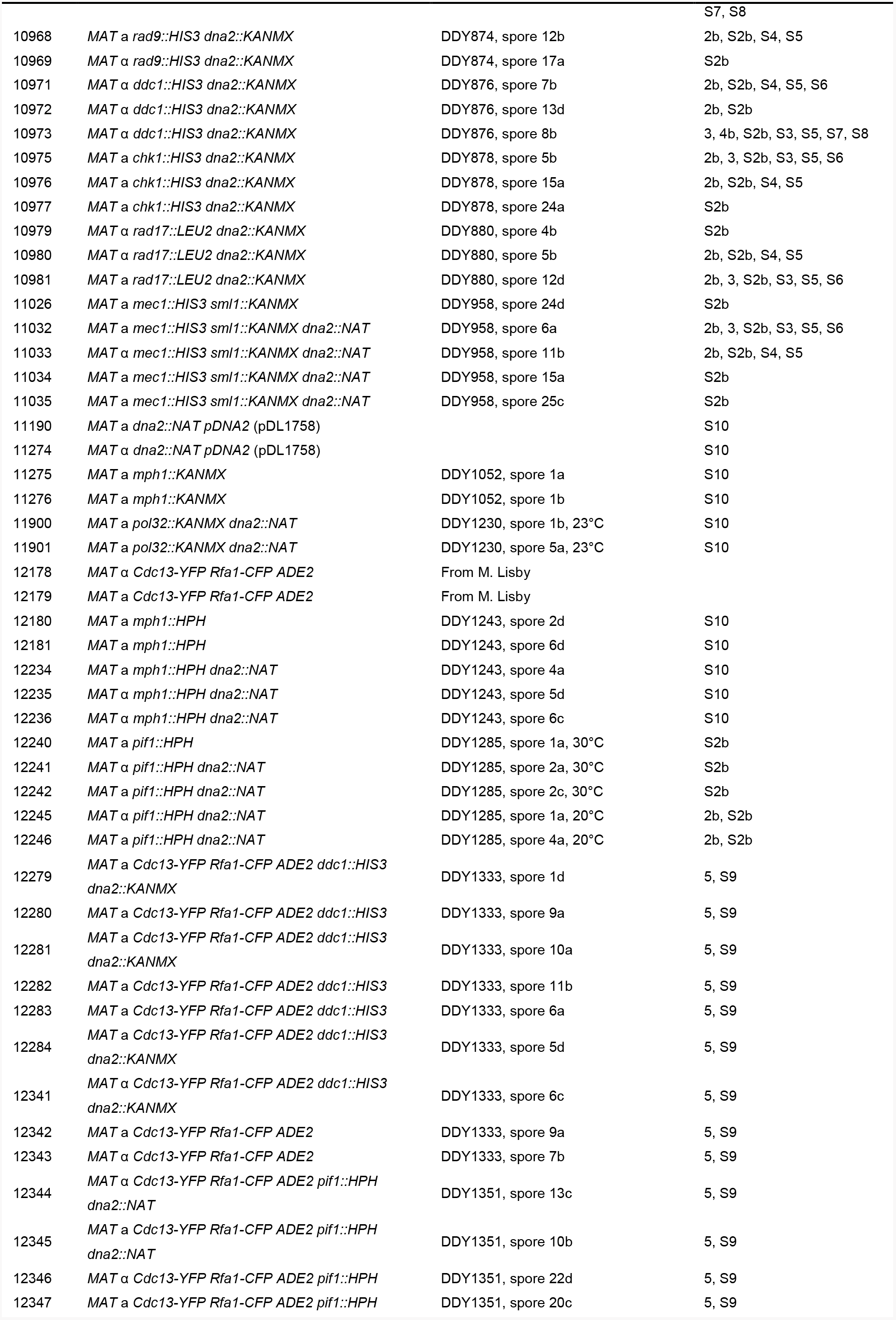
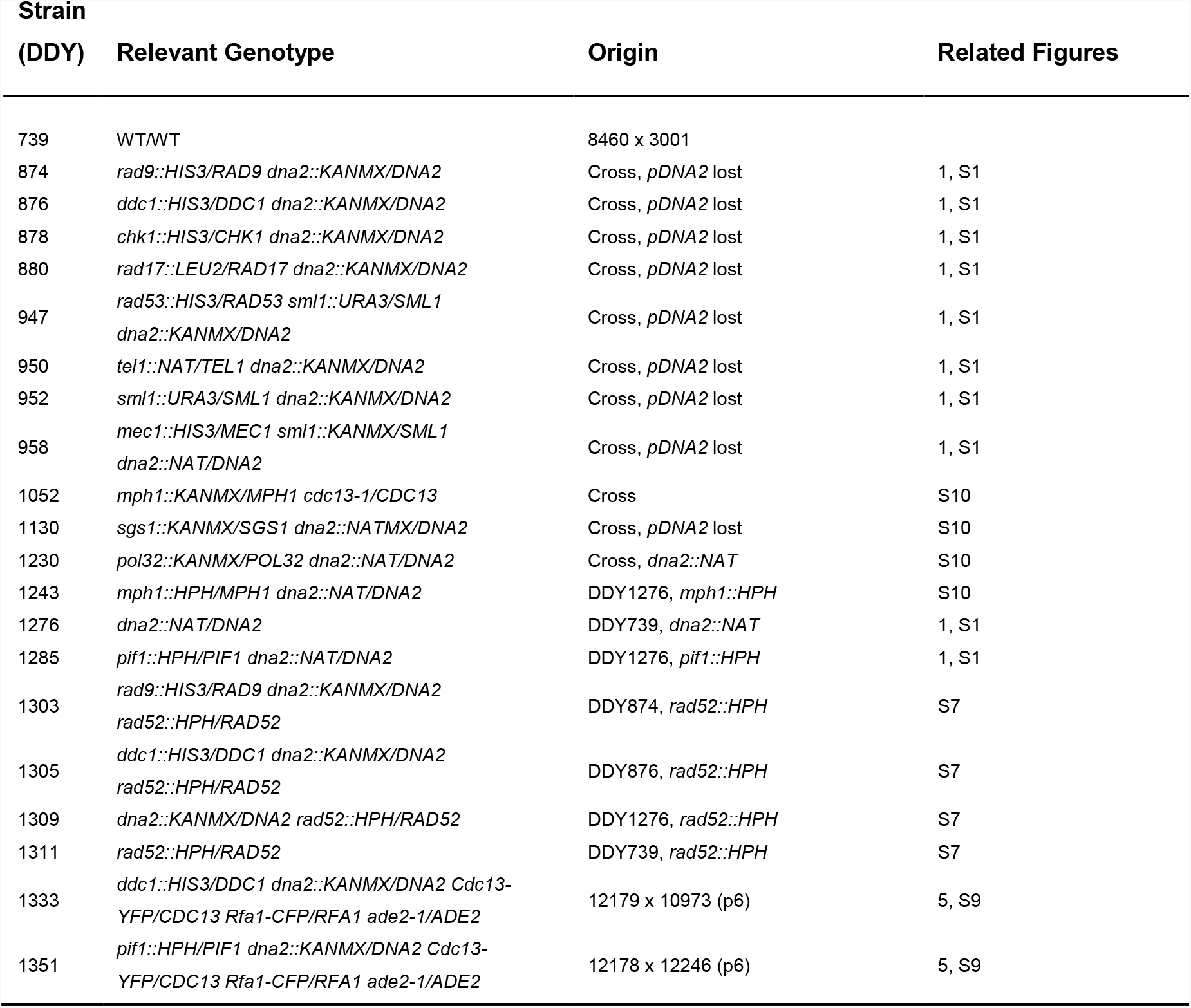
Yeast strains used. *S. cerevisiae* strains used are in the W303 genetic background (*ade2-1 can1-100 trp1-1 leu2-3,112 his3-11,15 ura3 GAL* + *psi*+ *ssd1-d2 RAD5*+), unless stated otherwise. Haploid strain numbers are prefixed DLY, diploids are DDY. Yeast strains are ordered by strain number.

**Supplementary Table 2.** List of plasmids used. Related figures indicated.

| **Plasmid Number** | **Details** | **Source (Related Figures)** |
| --- | --- | --- |
| pDL1367 | 2μ-*URA3* | pYEP24 (4, S8) |
| pDL1369 | 2μ-*URA3-DNA2* | Zhu *et al*, Cell 2008 19;134(6):981-94 (4b, S8) |
| pDL1371 | 2μ-*URA3-dna2-R1253Q* | Zhu *et al*, Cell 2008 19;134(6):981-94 (4b, S8) |
| pDL1373 | 2μ-*URA3-dna2-E675A* | Zhu *et al*, Cell 2008 19;134(6):981-94 (4b, S8) |
| pDL1539 | pRS314-*TRP1* | Pfander and Diffley, The EMBO Journal 2011 30: 4897-4907 (4b, S8) |
| pDL1544 | pRS314-*TRP1-DNA2* | Kumar and Burgers, Genes and Development 2013 27: 313-321 (4b, S8) |
| pDL1561 | pRS314-*TRP1-dna2-W128A, Y130A* | Kumar and Burgers, Genes and Development 2013 27: 313-321 (4, S8) |
| pDL1758 | pAG36::*CAN1-URA3-DNA2* | Goldstein, A. and J. McCusker, Yeast 1999 15(14):1541-53 (S10) |

**Figure S1.**
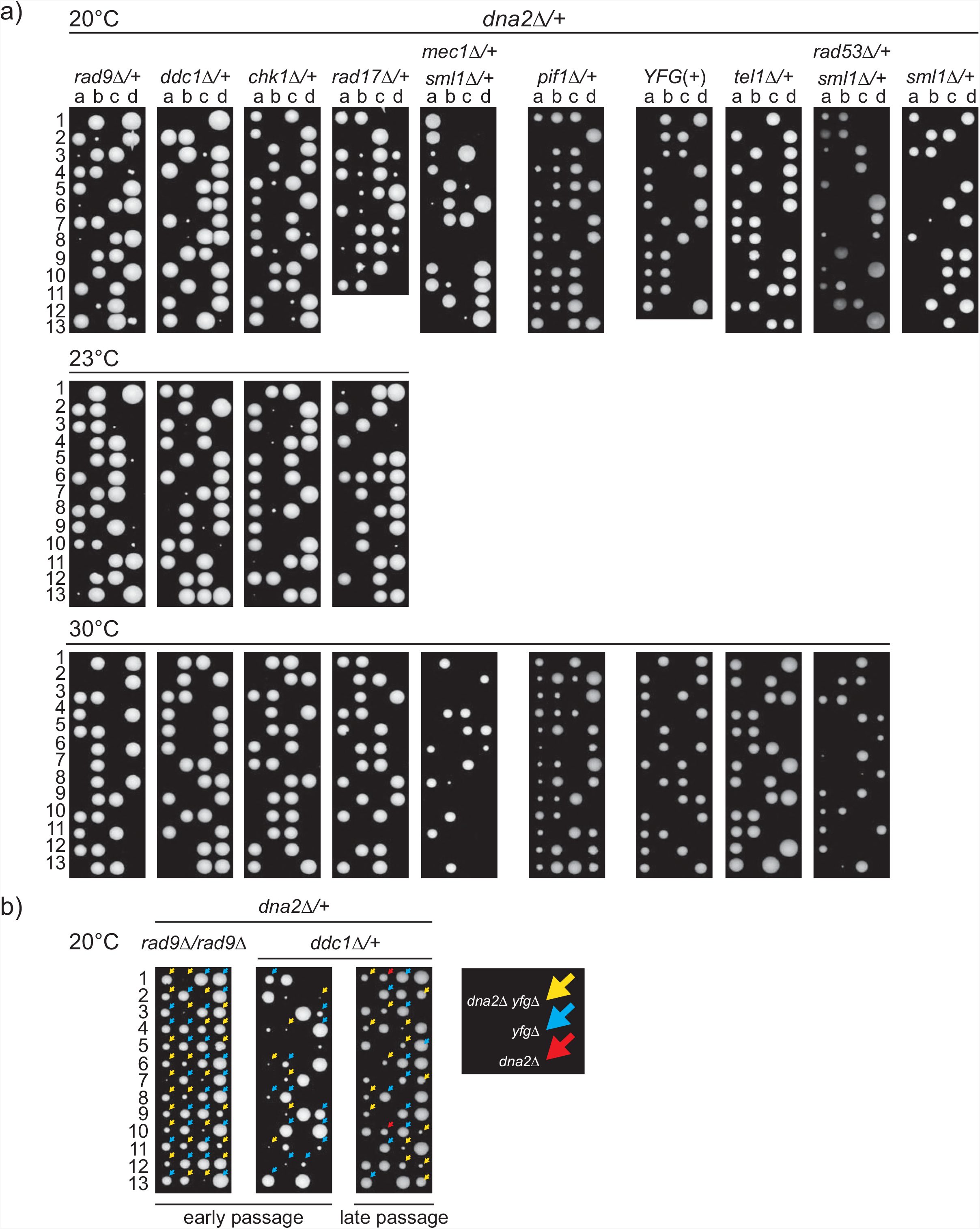
Checkpoint gene deletions affect *dna2*Δ viability. Germination plates as in Figure 1. a) Spores were germinated at 20,C, 23,C and 30°C for 10-11, 7 and 3-4 days before photographing, respectively. Strains were: DDY1285, DDY874, DDY876, DDY878, DDYBBO, DDY958, DDY950, DDY947, DDY952, DDY1276, strain details are in Suppl. Table 1. b) *dna2checkpoint* strains from passage 2 (early passage) or passage 6 (late passage) were crossed to *rad9*Δ or WT strain. *dna2*Δ strains are highlighted in red. Strains were: *dna2*Δ *rad9*Δ (DLY10967) x *rad9*Δ (DLY9593), *dna2*Δ *ddc1*Δ (DLY10973) x WT (DLY8460).

**Figure S2.**
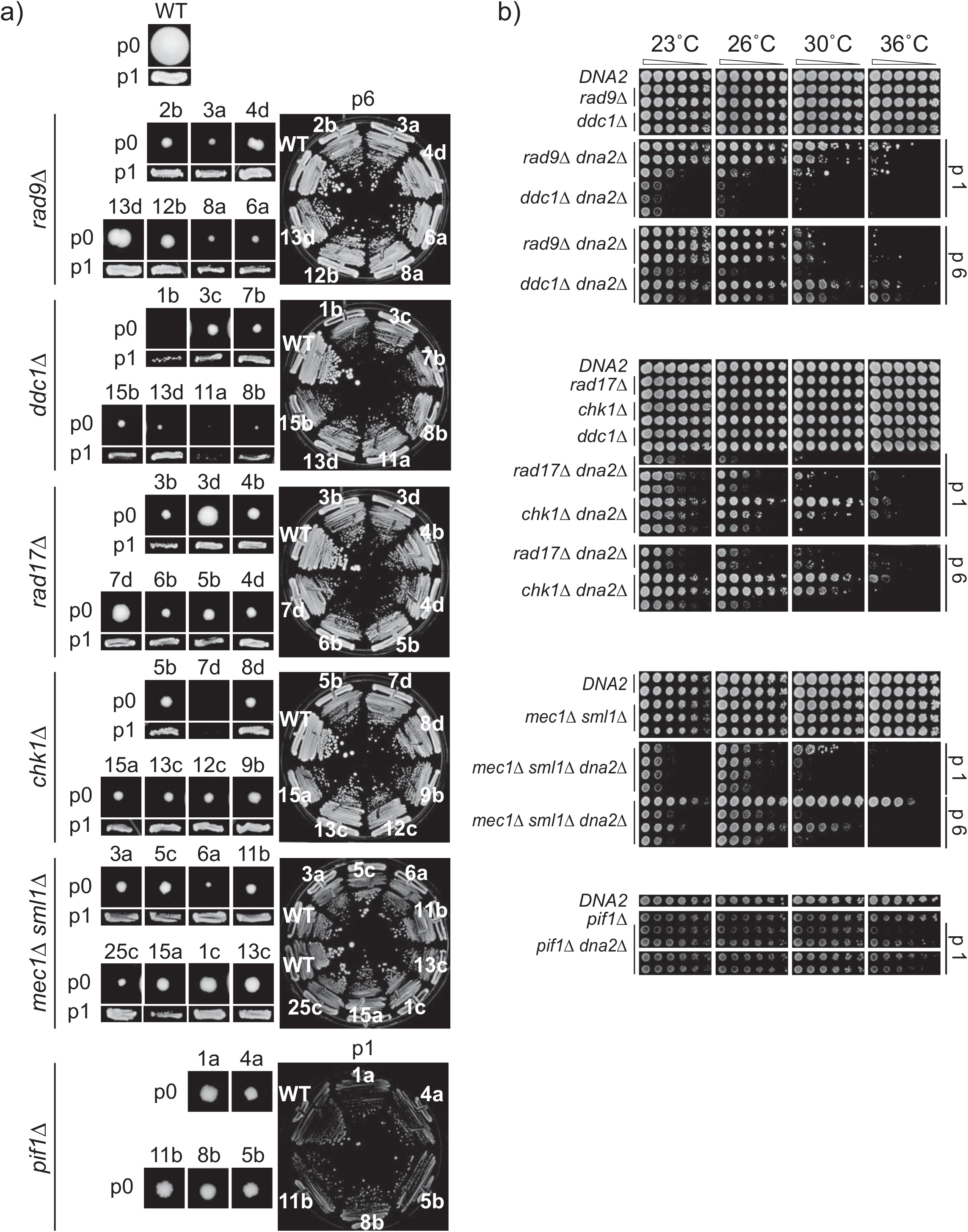
*dna2*Δ strains improve growth with passage, but remain temperature sensitive. a) Colony sizes from spores (passage 0), p1 and p6 of viable *dna2*Δ mutants, as in Figure 2a. b) Spot test assays as in Figure 2b. Strains were: WT (DLY3001, DLY8460), *rad9*Δ (DLY9593, DLY10818), *rad9*Δ (DLY7173, DLY7174), *rad9*Δ *dna2*Δ (DLY10967, DLY10968, DLY10969), *ddc1A dna2*Δ (DLY10971, DLY10972, DLY10973), *rad17*Δ (DLY7177, DLY7178), *chk1*Δ (DLY10536, DLY10537), *rad17A dna2*Δ (DLY10979, DLY10980, DLY10981), *chk1*Δ *dna2*Δ(DLY10975, DLY10976, DLY10977), *mec1*Δ *sml1*Δ (DLY1326, DLY6855, DLY11026), *mec1*Δ *sml1*Δ *dna2*Δ(DLY11032, DLY11033, DLY11034, DLY11035), *pif1*Δ (DLY12240), *pif1*Δ *dna2*Δ (DLY12241, DLY12242, DLY12245, DLY12246).

**Figure S3.**
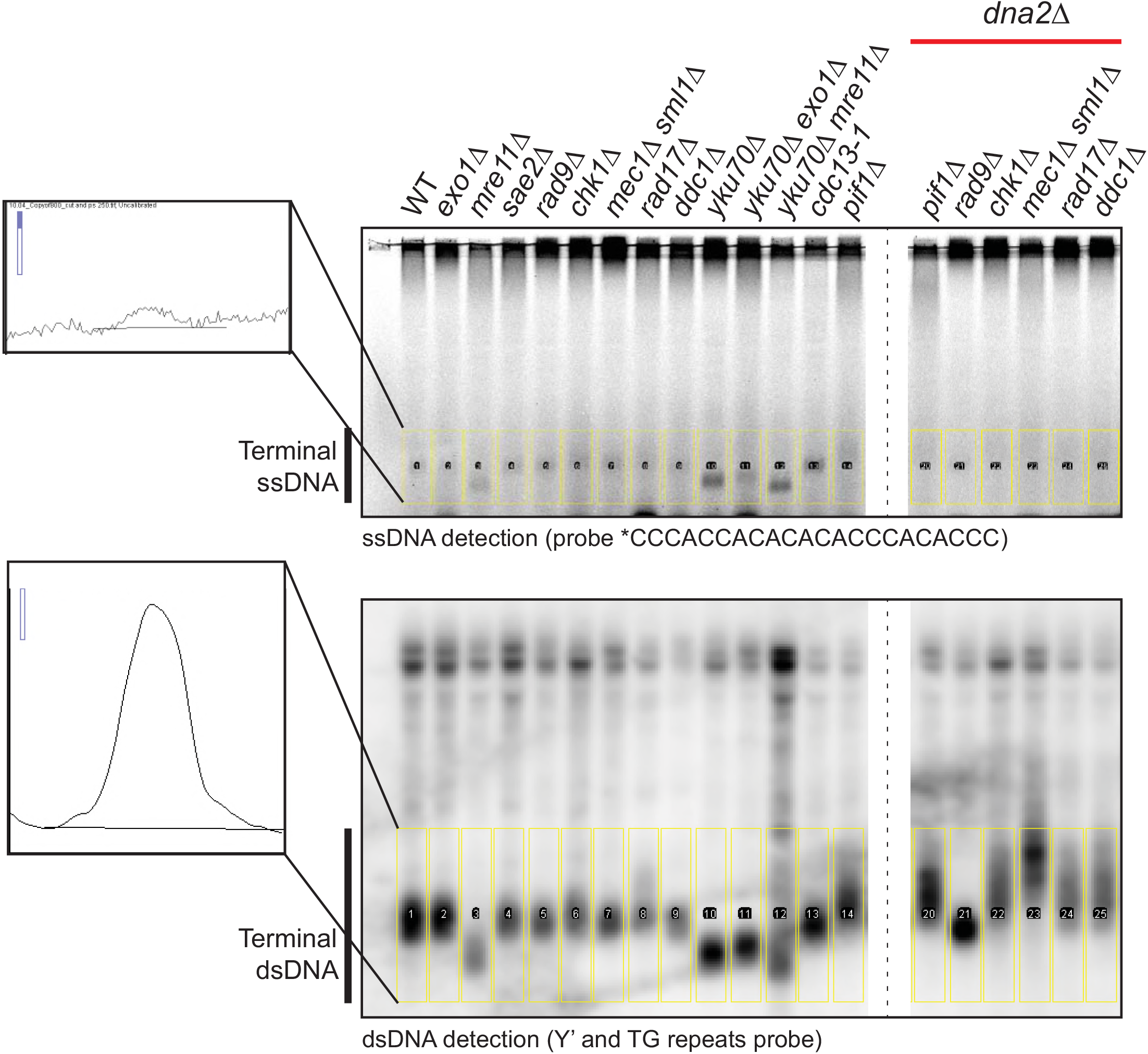
ssDNA and dsDNA quantification. Telomeric regions from Figure 3 were quantified using an ImageJ software, indicated by yellow rectangulars. ssDNA was quantified from in-gel assay (top panel), and dsDNA from a Southern blot (bottom panel). Graphs generated in ImageJ for a WT sample and the cut offs are shown.

**Figure S4.**
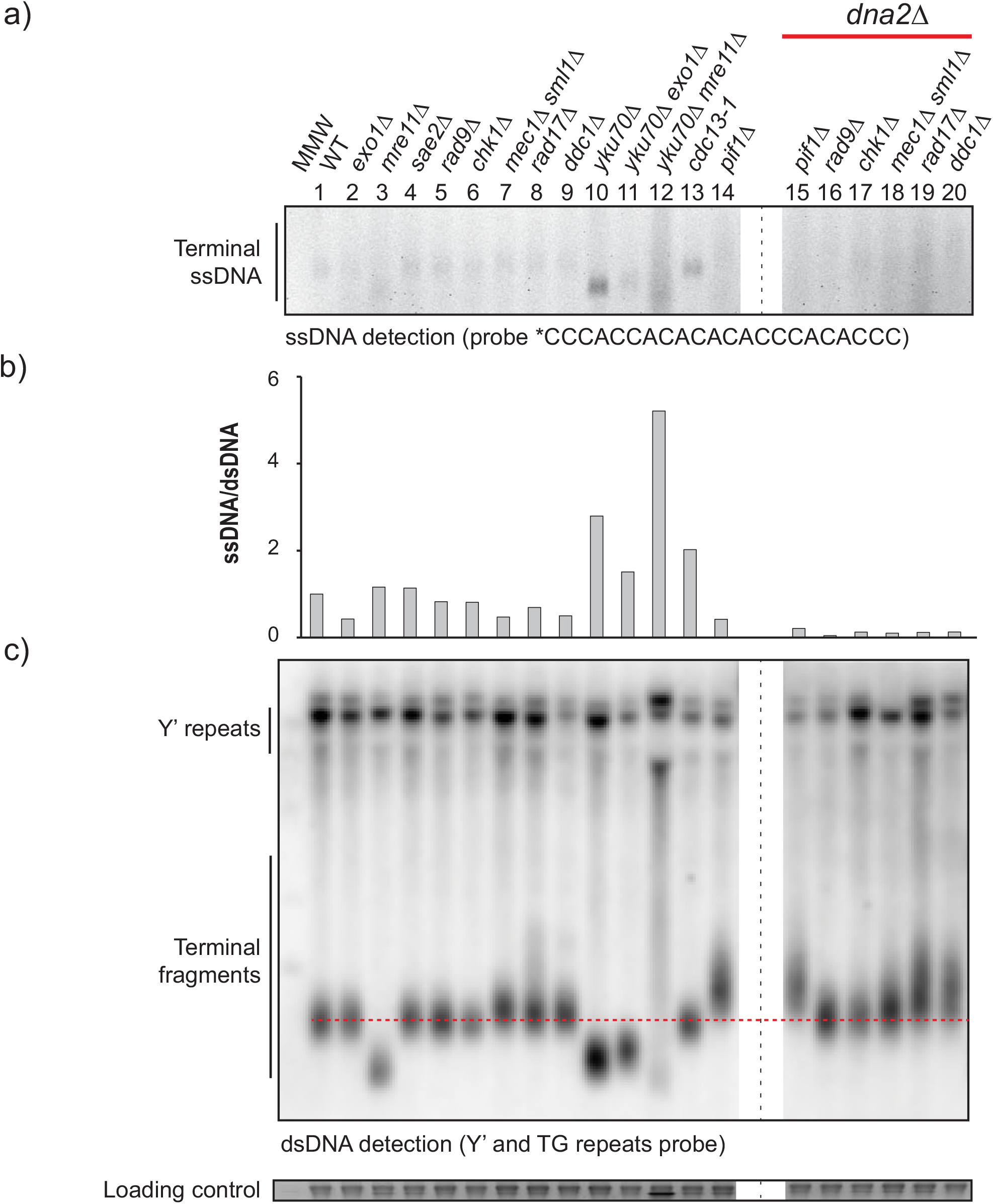
Telomeres of *dna2*Δ strains are abnormal and have low levels of ssDNA. a-c) In-gel and Southern blot analysis of independent strains of the same genotypes as shown in Figure 3. Strains were: WT (DLY8460), *exo1*Δ (DLY1273), *mre11*Δ (DLY4458), *sae2*Δ (DLY1578), *rad9*Δ (DLY658), *chk1*Δ (DLY1095), *mec1*Δ *sml1*Δ (DLY6855), *rad17*Δ (DLY7178), *ddc1*Δ (DLY7173), *yku70*Δ (DLY1412), *yku70*Δ *exo1*Δ (DLY1409), *yku70*Δ *mre11*Δ (DLY1846), *cdc13-1* (DLY1195), *pif1*Δ (DLY5394), *pif1*Δ *dna2*Δ (DLY4691), *rad9*Δ *dna2*Δ (DLY10968), *chk1*Δ *dna2*Δ (DLY10976), *mec1*Δ *sml1*Δ *dna2*Δ (DLY11033), *rad17*Δ *dna2*Δ (DLY10980), *ddc1*Δ *dna2*Δ (DLY10971). Strain details are in Suppl. Table 1.

**Figure S5.**
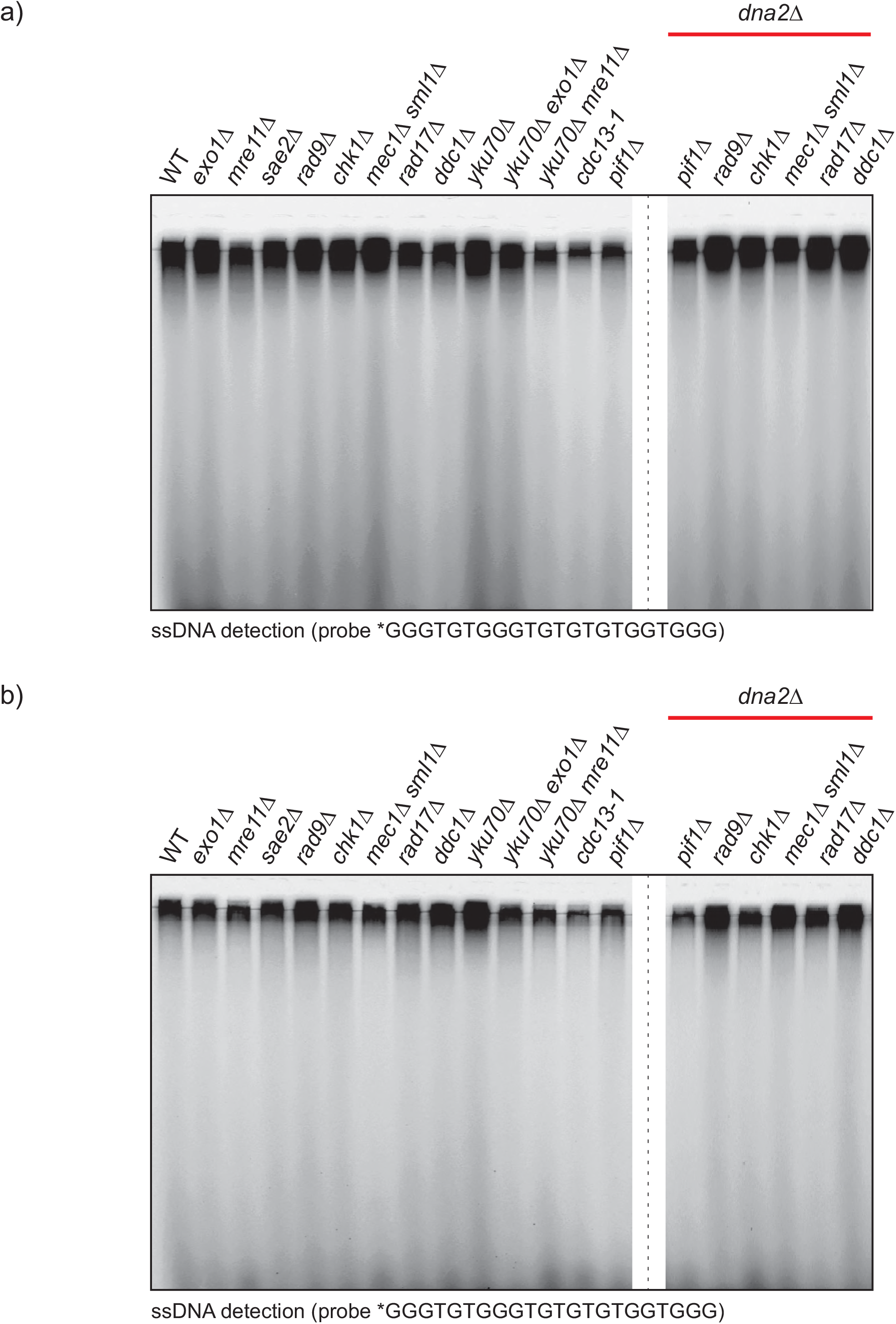
C-rich strand is not detected by in-gel assay. In-gel assays performed as in Figure 3, except that TG probe rather than AC probe was used. a) An in-gel assay on strains from Figure 3. b) An in-gel assay on strains from Figure S4.

**Figure S6.**
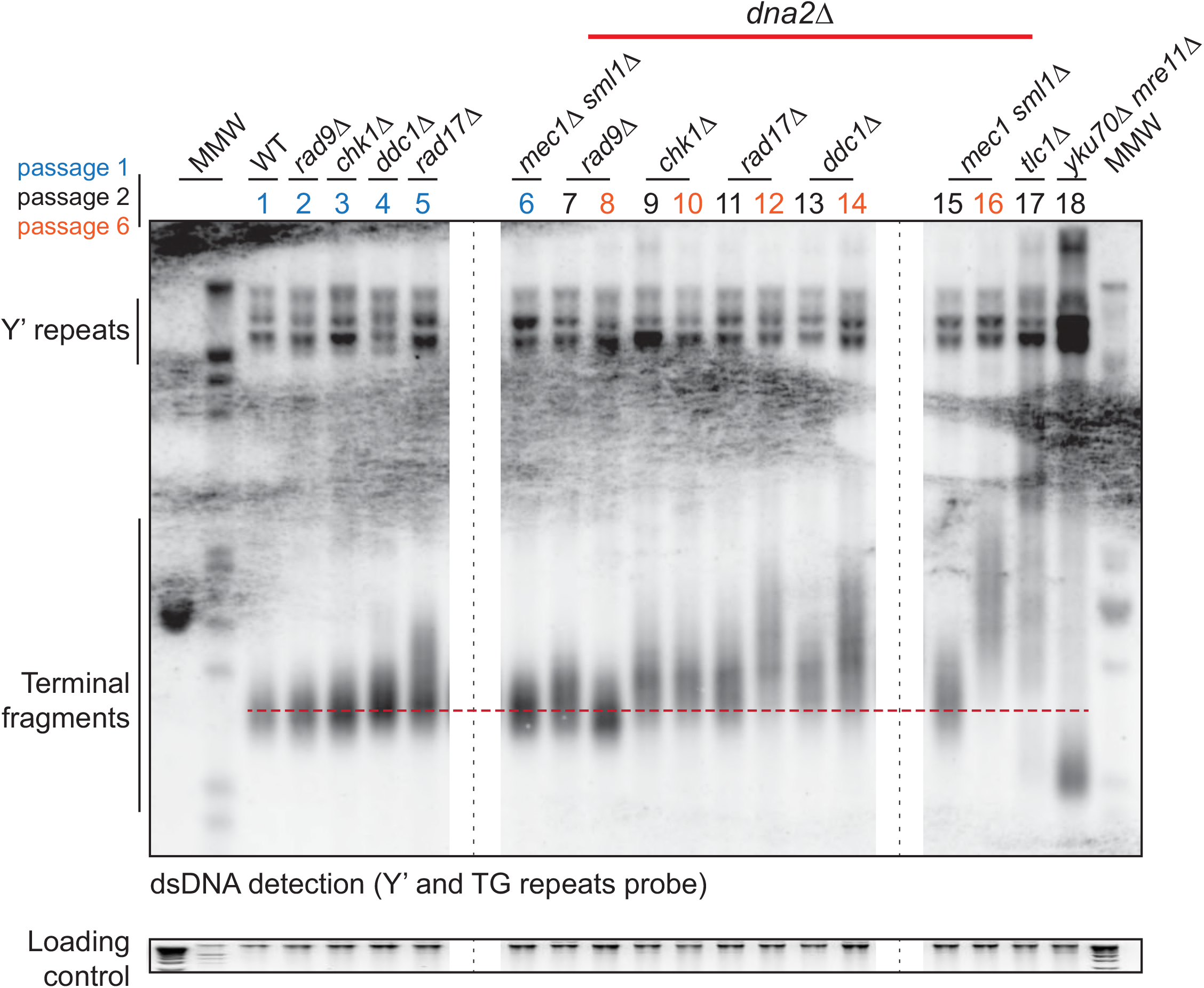
Telomeres of *dna2*Δ strains are abnormal. Southern blot performed as described previously (Maringele and Lydall, 2004). DNA was isolated from yeast strains grown in 2 mL YEPD until saturation at 23°C. Strains were: WT (DLY3001), *rad9*Δ (DLY9593), *chk1*Δ (DLY10536), *ddc1*Δ (DLY7173), *rad17*Δ (DLY7177), *mec1*Δ *sml1*Δ (DLY1326), *rad9*Δ *dna2*Δ (DLY10967), *chk1*Δ *dna2*Δ (DLY10975), *rad17*Δ *dna2*Δ (DLY10981), *ddc1*Δ *dna2*Δ (DLY10971), *mec1*Δ *sml1*Δ *dna2*Δ (DLY11032), *tlc1*Δ (DLY2147), *yku70*Δ *mre11*Δ (DLY1845). Strain details are in Suppl. Table 1.

**Figure S7.**
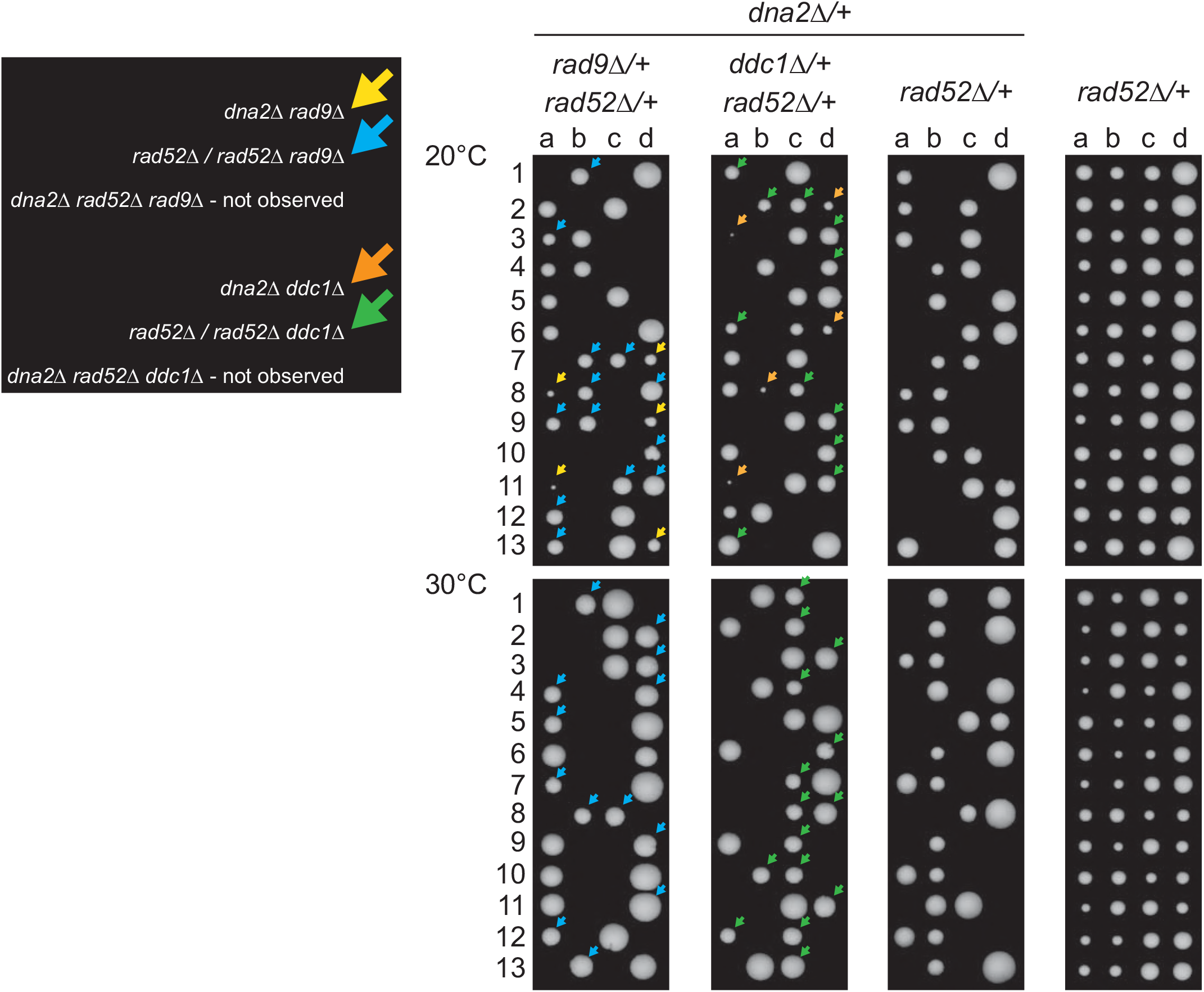
Suppression of *dna2*Δ is HR-dependent. RAD52 gene was deleted in DDY874 (DDY1303), DDY876 (DDY1305), DDY1276 (DDY1309) and DDY739 (DDY1311). Diploids were sporulated and germinated as in Figure 1. Arrows indicate colonies of appropriate genotypes, shown on the left. Strain details are in Suppl. Table 1.

**Figure S8.**
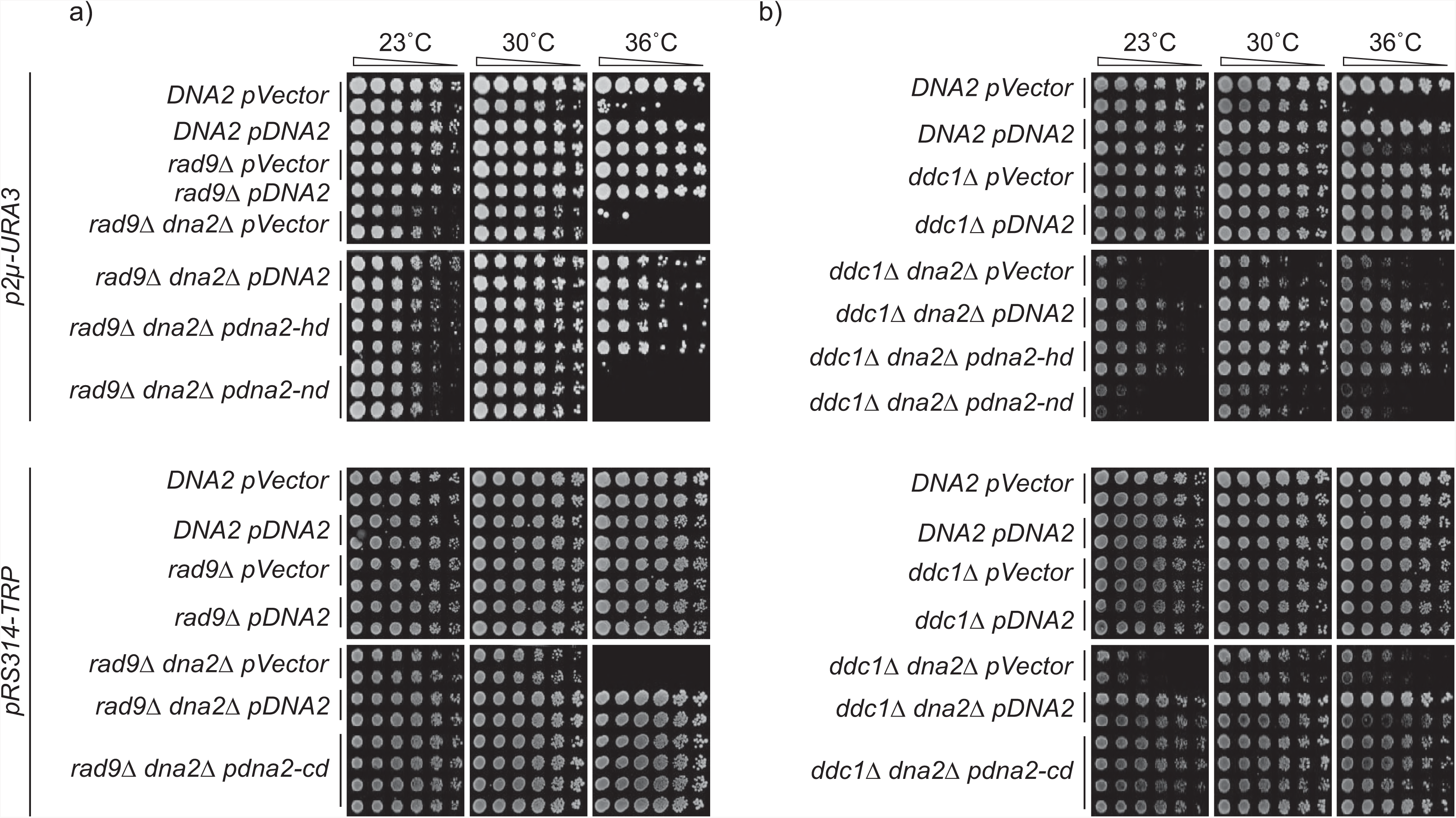
The nuclease domain of Dna2, but not helicase or checkpoint domains, confers viability of *dna2*Δ strains. Spot test assays performed as in Figure 4b. WT (DLY3001), *rad9*Δ (DLY9593), *rad9*Δ *dna2*Δ (DLY10967) (a) and WT (DLY3001), *ddc1*Δ (DLY8530), *ddc1*Δ *dna2*Δ (DLY10973) (b) strains carried plasmids: *pVector (2µ-J-URA3), pDNA2 (2µ-J-URA3-DNA2), pdna2-hd* (dna2-helicase dead, *2µ-J-URA3-dna2-R1253Q), pdna2-nd* (dna2-nuclease dead, *2µ-J-URA3-dna2-E675A), pVector* (pRS314-*TRP1), pDNA2* (pRS314-*TRP1-DNA2), pdna2-cd* (dna2-checkpoint-dead, *pRS314-TRP1-dna2-W128A,Y130A).*

**Figure S9.**
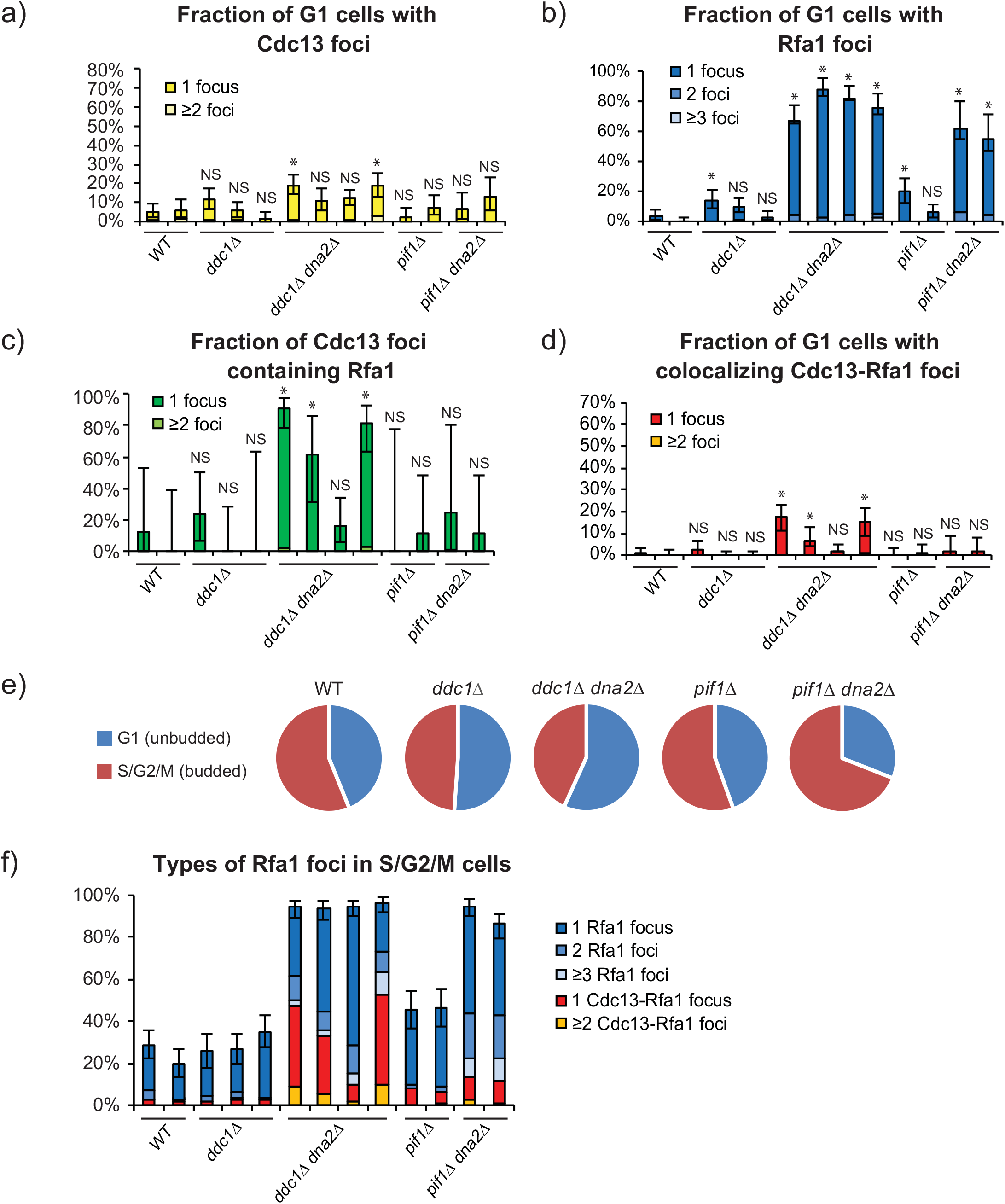
*dna2*Δ mutants accumulate CST and RPA, the ssDNA binding complexes. Percentages of Cdc13 foci, Rfa1 foci or colocalized Cdc13–Rfa1 foci in *dna2*Δ and control strains are shown. a) Percentage of unbudded (G1) cells with either Cdc13 foci only or Cdc13-Rfa1 foci. b) Percentage of unbudded cells with either Rfa1 foci only or Cdc13-Rfa1 foci. c) Percentage of Cdc13 foci that colocalize with Rfa1 foci. d) Percentage of unbudded cells with colocalizing Cdc13-Rfa1 foci. Error bars indicate 95 % confidence intervals (n=213-437, from two independent cultures of each strain). * - statistical significance (p<0.05) determined using Fisher’s exact test. NS - not significant. Strains are: WT (DLY12342, DLY12343), *ddc1*Δ (DLY12282, DLY12280, DLY12283), *ddc1*Δ *dna2*Δ (DLY12281, DLY12341, DLY12284, DLY12279), *pif1*Δ (DLY12346, DLY12347), *pif1*Δ *dna2*Δ (DLY12344, DLY12345). e) Cell cycle distribution of cells from panels a-d) was determined based cell morphology (budded (S/G2/M) versus unbudded (G1)). f) Separation of Rfa1 foci into telomeric (Cdc13 colocalizing) and non-telomeric (non-Cdc13 colocalizing) foci for S/G2/M cells. Strain details are in Suppl. Table 1.

**Figure S10.**
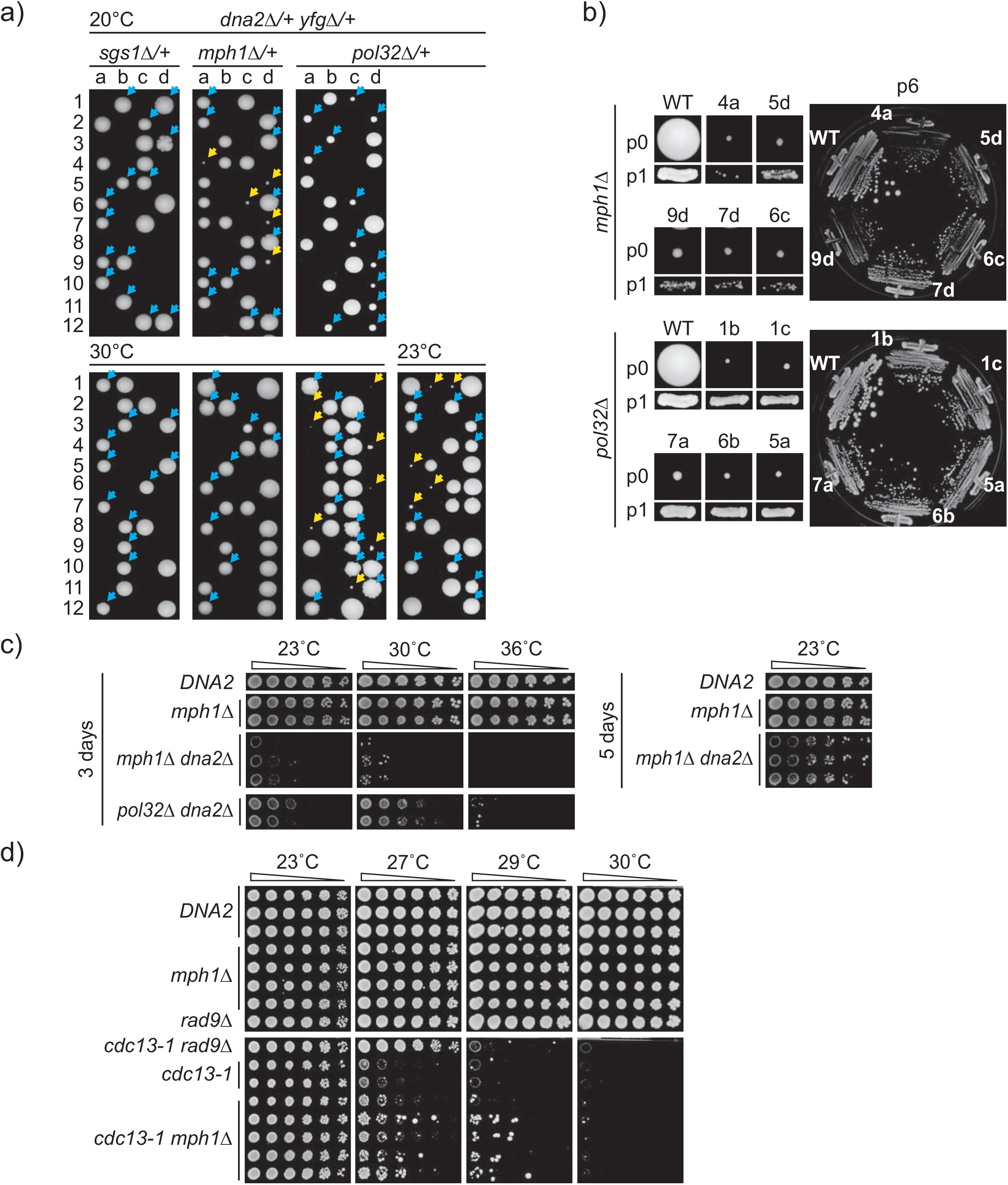
Deletion of *MPH1* and *POL32*, but not *SGS1*, suppress *dna2*Δ similarly to *checkpoint*Δ. a) Germination plates as in Figure 1. Spores were germinated at 20°C, 23°C and 30°C for 10, 7 and 3 days before photographing, respectively. Strains of *dna2*Δ *yfg*Δ background are indicated by yellow arrows, and strains of *yfg*Δ background are indicated by blue arrows. Strains were: DDY1130, DDY1243, DDY1230, strain details are in Suppl. Table 1. b-c) *dna2*Δ strains improve growth with passage, but remain temperature sensitive. b) Colony sizes from spores (passage 0), p1 and p6 of viable *dna2*Δ mutants, as in Figure 2a. c) Spot test assays as in Figure 2b. Strains were: WT (DLY3001), *mph1*Δ (DLY12180, DLY12181), *mph1*Δ *dna2*Δ (DLY12234, DLY12235, DLY12236), *pol32*Δ *dna2*Δ (DLY11900, DLY11901). d) *mph1*Δ suppresses growth defect of telomere defective *cdc13-1* strains. Spot test assays as in Figure 2b. Strains were: WT (DLY3001), *dna2*Δ pDNA2 (DLY11190, DLY11274), *mph1*Δ (DLY4282, DLY4283, DLY11275, DLY11276), *rad9*Δ (DLY9593), *rad9*Δ *cdc13-1* (DLY9585), *cdc13-1* (DLY1108, DLY1195), *mph1*Δ *cdc13-1* (DLY4106, DLY4107, DLY4108, DLY4285, DLY4286). Strain details are in Suppl. Table 1.

## References

Addinall, S. G., Downey, M., Yu, M., Zubko, M. K., Dewar, J., Leake, A., Hallinan, J., Shaw, O., James, K., Wilkinson, D. J., Wipat, A., Durocher, D. & Lydall, D. 2008. A genomewide suppressor and enhancer analysis of cdc13-1 reveals varied cellular processes influencing telomere capping in Saccharomyces cerevisiae. Genetics, 180, 2251–66.

Bae, S. H., Bae, K. H., Kim, J. A. & Seo, Y. S. 2001. RPA governs endonuclease switching during processing of Okazaki fragments in eukaryotes. Nature, 412, 456–61.

Balakrishnan, L. & Bambara, R. A. 2013. Okazaki fragment metabolism. Cold Spring Harb Perspect Biol, 5.

Bonetti, D., Martina, M., Falcettoni, M. & Longhese, M. P. 2013. Telomere-end processing: mechanisms and regulation. Chromosoma.

Bonetti, D., Villa, M., Gobbini, E., Cassani, C., Tedeschi, G. &Longhese, M. P. 2015. Escape of Sgs1 from Rad9 inhibition reduces the requirement for Sae2 and functional MRX in DNA end resection. EMBO Rep, 16,351–61.

Budd, M. E., Antoshechkin, I. A., Reis, C., Wold, B. J. & Campbell, J. L. 2011. Inviability of a DNA2 deletion mutant is due to the DNA damage checkpoint. Cell Cycle, 10, 1690–8.

Budd, M. E. & Campbell, J. L. 1997. A yeast replicative helicase, Dna2 helicase, interacts with yeast FEN-1 nuclease in carrying out its essential function. Mol Cell Biol, 17, 2136–42.

Budd, M. E. & Campbell, J. L. 2013. Dna2 is involved in CA strand resection and nascent lagging strand completion at native yeast telomeres. J Biol Chem, 288, 29414–29.

Budd, M. E., Choe, W. C. & Campbell, J. L. 1995. DNA2 encodes a DNA helicase essential for replication of eukaryotic chromosomes. J Biol Chem, 270, 26766–9.

Budd, M. E., Reis, C. C., Smith, S., Myung, K. & Campbell, J. L. 2006. Evidence suggesting that Pif1 helicase functions in DNA replication with the Dna2 helicase/nuclease and DNA polymerase delta. Mol Cell Biol, 26, 2490–500.

Budd, M. E., Tong, A. H. Y., Polaczek, P., Peng, X., Boone, C. & Campbell, J. 2005. A Network of Multi-tasking Proteins at the DNA replication Fork Preserves Genome Stability. PLoS Genetics,preprint, e61.

Bunting, S. F., Callen, E., Wong, N., Chen, H. T., Polato, F., Gunn, A., Bothmer, A., Feldhahn, N., Fernandez-Capetillo, O., Cao, L., Xu, X., Deng, C. X., Finkel, T., Nussenzweig, M., Stark, J. M. & Nussenzweig, A. 2010. 53BP1 inhibits homologous recombination in Brca1-deficient cells by blocking resection of DNA breaks. Cell, 141, 243–54.

Burgers, P. M. 2009. Polymerase dynamics at the eukaryotic DNA replication fork. J Biol Chem, 284, 4041–5.

Byrd, A. K. & Raney, K. D. 2015. A parallel quadruplex DNA is bound tightly but unfolded slowly by pif1 helicase. J Biol Chem, 290, 6482–94.

Ceccaldi, R., Sarangi, P. & D’Andrea, A. D. 2016. The Fanconi anaemia pathway: new players and new functions. Nat Rev Mol Cell Biol, 17, 337–49.

Cejka, P., Cannavo, E., Polaczek, P., Masuda-Sasa, T., Pokharel, S., Campbell, J. L. & Kowalczykowski, S. C. 2010. DNA end resection by Dna2-Sgs1-RPA and its stimulation by Top3-Rmi1 and Mre11-Rad50-Xrs2. Nature, 467, 112–6.

Chai, W., Zheng, L. & Shen, B. 2013. DNA2, a new player in telomere maintenance and tumor suppression. Cell Cycle, 12, 1985–6.

Choe, W., Budd, M., Imamura, O., Hoopes, L. & Campbell, J. L. 2002. Dynamic localization of an Okazaki fragment processing protein suggests a novel role in telomere replication. Mol Cell Biol, 22, 4202–17.

COSMIC. Available: http://cancer.sanger.ac.uk/cosmic.

Dewar, J. M. & Lydall, D. 2010. Pif1- and Exo1-dependent nucleases coordinate checkpoint activation following telomere uncapping. EMBO J, 29, 4020–34.

Dewar, J. M. & Lydall, D. 2012. Simple, non-radioactive measurement of single-stranded DNA at telomeric, sub-telomeric, and genomic loci in budding yeast. Methods Mol Biol, 920, 341–8.

Dominguez-Valentin, M., Therkildsen, C., Veerla, S., Jonsson, M., Bernstein, I., Borg, A. & Nilbert, M. 2013. Distinct gene expression signatures in lynch syndrome and familial colorectal cancer type x. PLoS One, 8, e71755.

Dubarry, M., Lawless, C., Banks, A. P., Cockell, S. & Lydall, D. 2015. Genetic Networks Required to Coordinate Chromosome Replication by DNA Polymerases alpha, delta, and epsilon in Saccharomyces cerevisiae. G3 (Bethesda), 5, 2187–97.

Duxin, J. P., Dao, B., Martinsson, P., Rajala, N., Guittat, L., Campbell, J. L., Spelbrink, J. N. & Stewart, S. A. 2009. Human Dna2 is a nuclear and mitochondrial DNA maintenance protein. Mol Cell Biol, 29, 4274–82.

Gerik, K. J., Li, X., Pautz, A. & Burgers, P. M. 1998. Characterization of the two small subunits of Saccharomyces cerevisiae DNA polymerase delta. J Biol Chem, 273, 19747–55.

Gilson, E. & Geli, V. 2007. How telomeres are replicated. Nat Rev Mol Cell Biol, 8, 825–38.

Heim, R. & Tsien, R. Y. 1996. Engineering green fluorescent protein for improved brightness, longer wavelengths and fluorescence resonance energy transfer. Curr. Biol., 6, 178–82.

Holstein, E. M., Clark, K. R. & Lydall, D. 2014. Interplay between nonsense-mediated mRNA decay and DNA damage response pathways reveals that Stn1 and Ten1 are the key CST telomere-cap components. Cell Rep, 7, 1259–69.

Holstein, E. M., Ngo, G., Lawless, C., Banks, P., Greetham, M., Wilkinson, D. & Lydall, D. 2017. Systematic Analysis of the DNA Damage Response Network in Telomere Defective Budding Yeast. G3 (Bethesda), 7, 2375–2389.

Hoopes, L. L. M., Budd, M., Choe, W., Weitao, T. & Campbell, J. L. 2002. Mutations in DNA Replication Genes Reduce Yeast Life Span. Molecular and Cellular Biology, 22, 4136–4146.

Hu, J., Sun, L., Shen, F., Chen, Y., Hua, Y., Liu, Y., Zhang, M., Hu, Y., Wang, Q., Xu, W., Sun, F., Ji, J., Murray, J. M., Carr, A. M. & Kong, D. 2012. The intra-S phase checkpoint targets Dna2 to prevent stalled replication forks from reversing. Cell, 149, 1221–32.

Iwabuchi, K., Basu, B. P., Kysela, B., Kurihara, T., Shibata, M., Guan, D., Cao, Y., Hamada, T., Imamura, K., Jeggo, P. A., Date, T. & Doherty, A. J. 2003. Potential role for 53BP1 in DNA end-joining repair through direct interaction with DNA. J Biol Chem, 278, 36487–95.

Jia, P. P., Junaid, M., Ma, Y. B., Ahmad, F., Jia, Y. F., Li, W. G. & Pei, D. S. 2017. Role of human DNA2 (hDNA2) as a potential target for cancer and other diseases: A systematic review. DNA Repair (Amst), 59, 9–19.

Kang, H. Y., Choi, E., Bae, S. H., Lee, K. H., Gim, B. S., Kim, H. D., Park, C., Macneill, S. A. & Seo, Y. S. 2000. Genetic analyses of Schizosaccharomyces pombe dna2(+) reveal that dna2 plays an essential role in Okazaki fragment metabolism. Genetics, 155, 1055–67.

Kang, Y. H., Kang, M. J., Kim, J. H., Lee, C. H., Cho, I. T., Hurwitz, J. & Seo, Y. S. 2009. The MPH1 gene of Saccharomyces cerevisiae functions in Okazaki fragment processing. J Biol Chem, 284, 10376–86.

Kao, H. I. & Bambara, R. A. 2003. The protein components and mechanism of eukaryotic Okazaki fragment maturation. Crit Rev Biochem Mol Biol, 38, 433–52.

Khadaroo, B., Teixeira, M. T., Luciano, P., Eckert-Boulet, N., Germann, S. M., Simon, M. N., Gallina, I., Abdallah, P., Gilson, E., Geli, V. & Lisby, M. 2009. The DNA damage response at eroded telomeres and tethering to the nuclear pore complex. Nat Cell Biol, 11, 980–7.

Kumar, S. & Burgers, P. M. 2013. Lagging strand maturation factor Dna2 is a component of the replication checkpoint initiation machinery. Genes Dev, 27, 313–21.

Kumar, S., Peng, X., Daley, J., Yang, L., Shen, J., Nguyen, N., Bae, G., Niu, H., Peng, Y., Hsieh, H. J., Wang, L., Rao, C., Stephan, C. C., Sung, P., Ira, G. & Peng, G. 2017. Inhibition of DNA2 nuclease as a therapeutic strategy targeting replication stress in cancer cells. Oncogenesis, 6, e319.

Lazzaro, F., Sapountzi, V., Granata, M., Pellicioli, A., Vaze, M., Haber, J. E., Plevani, P., Lydall, D. & Muzi-Falconi, M. 2008. Histone methyltransferase Dot1 and Rad9 inhibit single-stranded DNA accumulation at DSBs and uncapped telomeres. EMBO J, 27, 1502–12.

Lee, K. H., Lee, M. H., Lee, T. H., Han, J. W., Park, Y. J., Ahnn, J., Seo, Y. S. & Koo, H. S. 2003. Dna2 requirement for normal reproduction of Caenorhabditis elegans is temperature-dependent. Mol Cells, 15, 81–6.

Lee, S. E., Moore, J. K., Holmes, A., Umezu, K., Kolodner, R. D. & Haber, J. E. 1998. Saccharomyces Ku70, mre11/rad50 and RPA proteins regulate adaptation to G2/M arrest after DNA damage. Cell, 94, 399–409.

Levikova, M. & Cejka, P. 2015. The Saccharomyces cerevisiae Dna2 can function as a sole nuclease in the processing of Okazaki fragments in DNA replication. Nucleic Acids Res, 43, 7888–97.

Levikova, M., Klaue, D., Seidel, R. & Cejka, P. 2013. Nuclease activity of Saccharomyces cerevisiae Dna2 inhibits its potent DNA helicase activity. Proc Natl Acad Sci U S A, 110, E1992–2001.

Lin, W., Sampathi, S., Dai, H., Liu, C., Zhou, M., Hu, J., Huang, Q., Campbell, J., Shin-Ya, K., Zheng, L., Chai, W. & Shen, B. 2013. Mammalian DNA2 helicase/nuclease cleaves G-quadruplex DNA and is required for telomere integrity. EMBO J, 32, 1425–39.

Maestroni, L., Matmati, S. & Coulon, S. 2017. Solving the Telomere Replication Problem. Genes (Basel), 8.

Maga, G., Stucki, M., Spadari, S. & Hubscher, U. 2000. DNA polymerase switching: I. Replication factor C displaces DNA polymerase alpha prior to PCNA loading. J Mol Biol, 295, 791–801.

Maga, G., Villani, G., Tillement, V., Stucki, M., Locatelli, G. A., Frouin, I., Spadari, S. & Hubscher, U. 2001. Okazaki fragment processing: modulation of the strand displacement activity of DNA polymerase delta by the concerted action of replication protein A, proliferating cell nuclear antigen, and flap endonuclease-1. Proc Natl Acad Sci U S A, 98, 14298–303.

Maringele, L. & Lydall, D. 2002. EXO1-dependent single-stranded DNA at telomeres activates subsets of DNA damage and spindle checkpoint pathways in budding yeast yku70Delta mutants. Genes Dev, 16, 1919–33.

Masuda-Sasa, T., Polaczek, P., Peng, X. P., Chen, L. & Campbell, J. L. 2008. Processing of G4 DNA by Dna2 helicase/nuclease and replication protein A (RPA) provides insights into the mechanism of Dna2/RPA substrate recognition. J Biol Chem, 283, 24359–73.

Mimitou, E. P. & Symington, L. S. 2008. Sae2, Exo1 and Sgs1 collaborate in DNA double-strand break processing. Nature, 455, 770–4.

Myler, L. R., Gallardo, I. F., Zhou, Y., Gong, F., Yang, S. H., Wold, M. S., Miller, K. M., Paull, T. T. & Finkelstein, I. J. 2016. Single-molecule imaging reveals the mechanism of Exo1 regulation by single-stranded DNA binding proteins. Proc Natl Acad Sci U S A, 113, E1170–9.

Navadgi-Patil, V. M. & Burgers, P. M. 2009a. A tale of two tails: activation of DNA damage checkpoint kinase Mec1/ATR by the 9-1-1 clamp and by Dpb11/TopBP1. DNA Repair (Amst), 8, 996–1003.

Navadgi-Patil, V. M. & Burgers, P. M. 2009b. The unstructured C-terminal tail of the 9-1-1 clamp subunit Ddc1 activates Mec1/ATR via two distinct mechanisms. Mol Cell, 36, 743–53.

Ngo, G. H., Balakrishnan, L., Dubarry, M., Campbell, J. L. & Lydall, D. 2014. The 9-1-1 checkpoint clamp stimulates DNA resection by Dna2-Sgs1 and Exo1. Nucleic Acids Res, 42, 10516–28.

Ngo, G. H. & Lydall, D. 2015. The 9-1-1 checkpoint clamp coordinates resection at DNA double strand breaks. Nucleic Acids Res, 43, 5017–32.

Nugent, C. I., Hughes, T. R., Lue, N. F. & Lundblad, V. 1996. Cdc13p: a single-strand telomeric DNA-binding protein with a dual role in yeast telomere maintenance. Science, 274, 249–52.

Ormo, M., Cubitt, A. B., Kallio, K., Gross, L. A., Tsien, R. Y. & Remington, S. J. 1996. Crystal structure of the Aequorea victoria green fluorescent protein [see comments]. Science, 273, 1392–5.

Paeschke, K., Bochman, M. L., Garcia, P. D., Cejka, P., Friedman, K. L., Kowalczykowski, S. C. & Zakian, V. A. 2013. Pif1 family helicases suppress genome instability at G-quadruplex motifs. Nature, 497, 458–62.

Parenteau, J. & Wellinger, R. J. 1999. Accumulation of single-stranded DNA and destabilization of telomeric repeats in yeast mutant strains carrying a deletion of RAD27. Mol Cell Biol, 19, 4143–52.

Peng, G., Dai, H., Zhang, W., Hsieh, H. J., Pan, M. R., Park, Y. Y., Tsai, R. Y., Bedrosian, I., Lee, J. S., Ira, G. & Lin, S. Y. 2012. Human nuclease/helicase DNA2 alleviates replication stress by promoting DNA end resection. Cancer Res, 72, 2802–13.

Phillips, J. A., Chan, A., Paeschke, K. & Zakian, V. A. 2015. The pif1 helicase, a negative regulator of telomerase, acts preferentially at long telomeres. PLoS Genet, 11, e1005186.

Pike, J. E., Burgers, P. M., Campbell, J. L. & Bambara, R. A. 2009. Pif1 helicase lengthens some Okazaki fragment flaps necessitating Dna2 nuclease/helicase action in the two-nuclease processing pathway. J Biol Chem, 284, 25170–80.

Podust, V. N., Podust, L. M., Muller, F. & Hubscher, U. 1995. DNA polymerase delta holoenzyme: action on single-stranded DNA and on double-stranded DNA in the presence of replicative DNA helicases. Biochemistry, 34, 5003–10.

Puddu, F., Granata, M., Di Nola, L., Balestrini, A., Piergiovanni, G., Lazzaro, F., Giannattasio, M., Plevani, P. & Muzi-Falconi, M. 2008. Phosphorylation of the budding yeast 9-1-1 complex is required for Dpb11 function in the full activation of the UV-induced DNA damage checkpoint. Mol Cell Biol, 28, 4782–93.

Rossi, M. L. & Bambara, R. A. 2006. Reconstituted Okazaki fragment processing indicates two pathways of primer removal. J Biol Chem, 281, 26051–61.

Schulz, V. P. & Zakian, V. A. 1994. The saccharomyces PIF1 DNA helicase inhibits telomere elongation and de novo telomere formation. Cell, 76, 145–55.

Sherman, F., Fink, G. R. & Hicks, J. B. 1986. Methods in Yeast Genetics,Cold Spring Harbor, NY, Cold Spring Harbor Laboratory.

Shim, E. Y., Chung, W. H., Nicolette, M. L., Zhang, Y., Davis, M., Zhu, Z., Paull, T. T., Ira, G. & Lee, S. E. 2010. Saccharomyces cerevisiae Mre11/Rad50/Xrs2 and Ku proteins regulate association of Exo1 and Dna2 with DNA breaks. EMBO J, 29, 3370–80.

Silva, S., Gallina, I., Eckert-Boulet, N. & Lisby, M. 2012. Live Cell Microscopy of DNA Damage Response in Saccharomyces cerevisiae. Methods Mol Biol, 920, 433–43.

Soudet, J., Jolivet, P. & Teixeira, M. T. 2014. Elucidation of the DNA end-replication problem in Saccharomyces cerevisiae. Mol Cell, 53, 954–64.

Stewart, J. A., Miller, A. S., Campbell, J. L. & Bambara, R. A. 2008. Dynamic removal of replication protein A by Dna2 facilitates primer cleavage during Okazaki fragment processing in Saccharomyces cerevisiae. J Biol Chem, 283, 31356–65.

Stith, C. M., Sterling, J., Resnick, M. A., Gordenin, D. A. & Burgers, P. M. 2008. Flexibility of eukaryotic Okazaki fragment maturation through regulated strand displacement synthesis. J Biol Chem, 283, 34129–40.

Strauss, C., Kornowski, M., Benvenisty, A., Shahar, A., Masury, H., Ben-Porath, I., Ravid, T., Arbel-Eden, A. & Goldberg, M. 2014. The DNA2 nuclease/helicase is an estrogen-dependent gene mutated in breast and ovarian cancers. Oncotarget, 5, 9396–409.

Sugiyama, T., Zaitseva, E. M. & Kowalczykowski, S. C. 1997. A single-stranded DNA-binding protein is needed for efficient presynaptic complex formation by the Saccharomyces cerevisiae Rad51 protein. J Biol Chem, 272, 7940–5.

Tomita, K., Kibe, T., Kang, H. Y., Seo, Y. S., Uritani, M., Ushimaru, T. & Ueno, M. 2004. Fission yeast Dna2 is required for generation of the telomeric single-strand overhang. Mol Cell Biol, 24, 9557–67.

Waga, S. & Stillman, B. 1998. The DNA replication fork in eukaryotic cells. Annu Rev Biochem, 67, 721–51.

Weinert, T. A., Kiser, G. L. & Hartwell, L. H. 1994. Mitotic checkpoint genes in budding yeast and the dependence of mitosis on DNA replication and repair. Genes Dev, 8, 652–65.

Weitao, T., Budd, M. & Campbell, J. L. 2003. Evidence that yeast SGS1, DNA2, SRS2, and FOB1 interact to maintain rDNA stability. Mutat Res, 532, 157–72.

Wellinger, R. J. & Zakian, V. A. 2012. Everything you ever wanted to know about Saccharomyces cerevisiae telomeres: beginning to end. Genetics, 191, 1073–105.

Wu, P., Takai, H. & DE Lange, T. 2012. Telomeric 3′ overhangs derive from resection by Exo1 and Apollo and fill-in by POT1b-associated CST. Cell, 150, 39–52.

Zhu, Z., Chung, W. H., Shim, E. Y., Lee, S. E. & Ira, G. 2008. Sgs1 helicase and two nucleases Dna2 and Exo1 resect DNA double-strand break ends. Cell, 134, 981–94.

Zou, L. & Elledge, S. J. 2003. Sensing DNA damage through ATRIP recognition of RPA-ssDNA complexes. Science, 300, 1542–8.

